# De Novo Design of Protein Switches with Diffusion-Based Ensemble Sampling

**DOI:** 10.64898/2026.07.20.739027

**Authors:** Alireza Omidi, Jiajun He, Jennifer M. Bui, Jörg Gsponer, Saifuddin Syed

## Abstract

Protein switches are proteins that can respond to biochemical stimuli by rearranging their structural elements, essential for cells to transduce signals. The de novo design of such proteins requires amino acid sequences whose energy landscapes support multiple stimulus-dependent conformations, yet most current de novo protein design pipelines are optimized for single stable structures. Existing multi-state inverse-folding methods can design sequences compatible with multiple backbones, but they assume that suitable backbone ensembles are already available, often requiring expert knowledge. We introduce Diff-Switch, a framework for sampling switch-like backbone ensembles from pretrained protein diffusion models. Given a reference backbone structure and domain decomposition, our method preserves local domain geometry while encouraging diversity in global domain arrangements along user-specified collective variables, inspired by metadynamics. We implement this objective through a controlled diffusion sampler with reward-tilting for local similarity between the ensemble members and history-dependent bias in collective-variable space to avoid repeated sampling of the same global arrangement. The resulting ensembles provide candidate conformational states for downstream multi-state inverse folding. Across our evaluation set of 20 diverse proteins, using conformations from these generated ensembles improves the success rate of finding switch-compatible sequences over baseline sampling. We further apply the method to a real-world protein switch design task and characterize the resulting designs.

## 1 Introduction

Proteins are central workhorses of cellular systems, involved in nearly all biological processes. Their function is tightly linked to their three-dimensional structures, i.e., the conformations they adopt in the cell. The development of machine-learning-based tools such as AlphaFold2 [Jumper et al., 2021] and AlphaFold3 [Abramson et al., 2024] has not only revolutionized our ability to predict protein structures but also enabled new approaches for computational protein design with tailored properties.

Despite these advances, a fundamental limitation remains: current approaches largely ignore or only partially capture protein dynamics. Conformational flexibility is often central to protein function [Praetorius et al., 2023]. Many proteins, including enzymes [Stiller et al., 2022], receptors [Bisello et al., 2002], and ion channels [Woll et al., 2021], operate by transitioning between distinct conformations. Proteins containing structural “switches”, elements that change conformation in response to specific stimuli, play key roles in regulation and signal transduction [Alberstein et al., 2022]. These switches can be triggered by ligand binding, post-translational modifications, or environmental changes, and span structural responses ranging from local rearrangements to large-scale domain motions or complete fold switching.

Designing protein switches is of high interest. However, their de novo design remains markedly more difficult than designing static structures. The challenge is inherently multi-objective: a single sequence must encode multiple stable conformations, enable transitions between them, and couple these transitions to a specific trigger [Alberstein et al., 2022]. In contrast, existing protein design pipelines are implicitly biased toward sequences with a single dominant free-energy minimum [Watson et al., 2023, Ingraham et al., 2023, Geffner et al., 2025, Lin et al., 2024, Yim et al., 2023, Zambaldi et al., 2024, Team et al., 2025, Stark et al., 2025]. These pipelines generate backbone structures, design sequences via inverse folding [Dauparas et al., 2022, Hsu et al., 2022, Ingraham et al., 2023, Shuai et al., 2025], and filter candidates using structure prediction models [Jumper et al., 2021, Abramson et al., 2024, Discovery et al., 2024, Passaro et al., 2025, Team et al., 2026], but fundamentally optimize for single-state stability rather than multi-state behavior.

Recent multi-state inverse-folding methods partially address this limitation by conditioning on multiple backbone conformations, e.g. ProteinMPNN-MSD [Dauparas et al., 2022] and DynamicMPNN [Abrudan et al., 2026]). However, their success critically depends on the availability of appropriate backbone ensembles. In practice, such ensembles are not systematically generated but instead constructed through manual design, domain expertise, or physics-based relaxation, making them difficult to obtain and scale [Langan et al., 2019].

On the other hand, through large-scale training, protein backbone diffusion models have learned rich knowledge of the protein conformation space. We leverage this knowledge in the design of protein switches. To this end we introduce Diff-Switch, a method for sampling diverse backbone ensembles from protein diffusion models while explicitly controlling switch reaction coordinates. By directly shaping the structural ensemble provided to multi-state inverse-folding methods, our approach enables the design of sequences more reliably realizing functional protein switches. Additionally, we apply our approach to a real-world design problem: the design of phosphorylation-induced protein switches.

## 2 Background

### 2.1 Protein switches

A protein consists of a chain of *L* amino acids bound by consecutive peptide bonds. Each amino acid is a molecule containing multiple atoms that can be categorised into backbone or side-chain atoms. Although amino acids differ in their side-chains, the backbone atoms are common to all 20 standard amino acids: the four backbone atoms N, C_*α*_, C, O can each be parameterised by their coordinates in ℝ^3^ and form the core of the protein chain. A protein backbone structure *x*, also known as a backbone conformation, then resides in X = ℝ^*L×*4*×*3^. A protein is fully described only when the sequence of its amino-acid side-chains, the protein sequence *s* = (*s*_1_, *s*_2_, …, *s*_*L*_), is also known.

While most protein sequences are thought to adopt one stable fold, which can be represented by a single conformation, this picture fails for protein switches, whose sequences encode at least two stable conformations and transition between them in response to a biochemical stimulus, e.g. binding to a small molecule. Formally, a protein sequence *s* is a switch if there exist backbone structures *x, x*^*′*^ ∈X that it can adopt, together with a reversible switch mechanism for the protein to go from one conformation to the other. See (A) and (B) in Figure 1 for a visualization of a protein switch and an example. For an in-depth review of protein switches and their design principles, we refer the reader to Alberstein et al. [2022].

**Figure 1.**
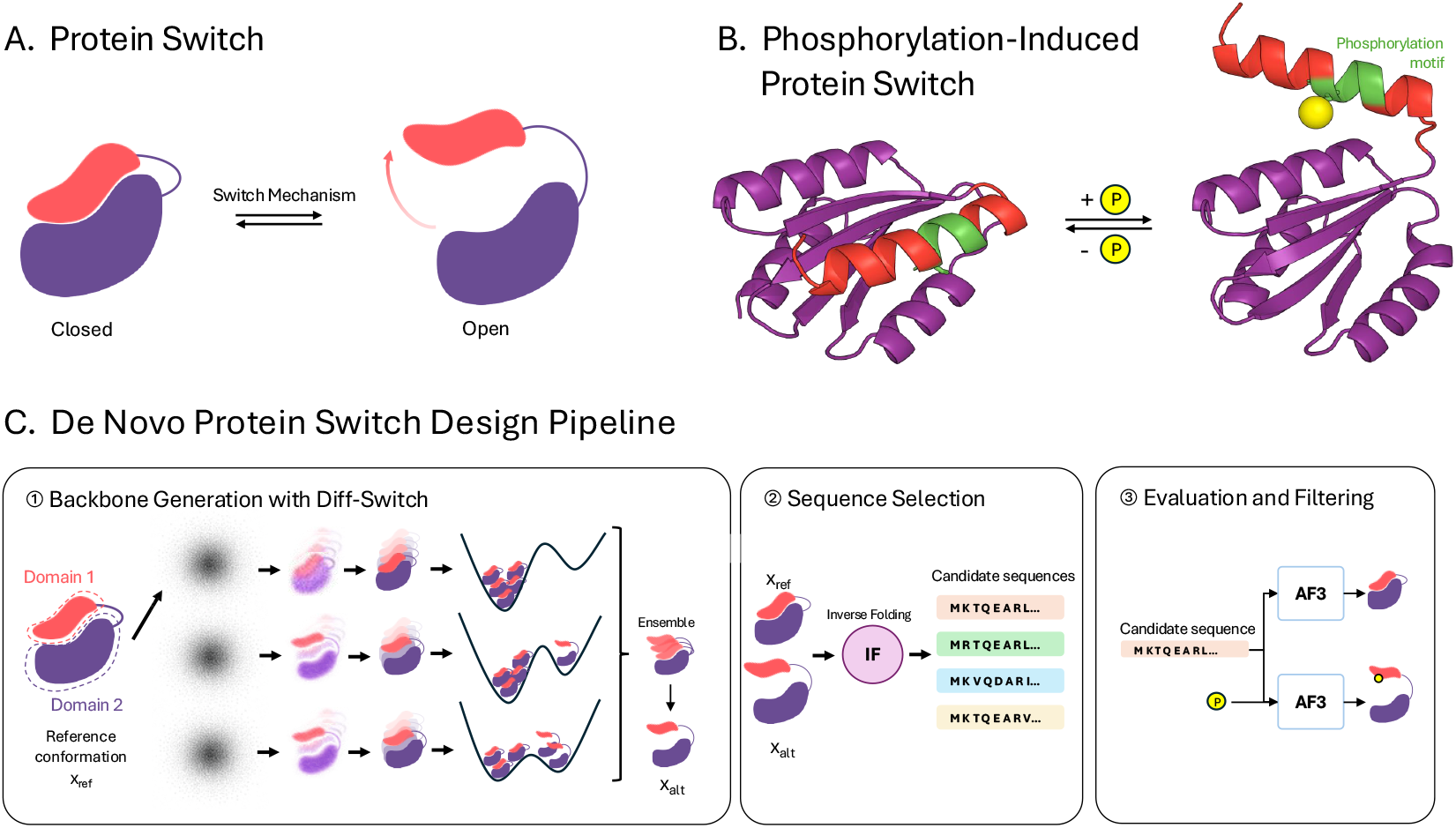
Protein switch design problem and overview of our Diff-Switch applied to phosphorylation-induced protein switches (Section 5.2). **(A)** Schematic of a protein switch: a single amino-acid sequence that adopts two stable conformations (closed and open) and transitions between them via a reversible switch mechanism. **(B)** Visualisation of a phosphorylation-induced protein switch [Buckley et al., 2025] generated by our de novo protein switch design pipeline. Addition or removal of the phosphate (yellow) on the switch helix containing the phosphorylation motif (green) opens or closes the protein. See Section 5.2 for details. **(C)** The three stages of our de novo protein switch design pipeline, applied here to phosphorylation-induced switching. The input is a reference backbone *x*_ref_ with annotated domains (red, purple). (1) Diff-Switch (Section 3.3) combines reward-tilted diffusion with well-tempered metadynamics to produce an ensemble ℰ of backbones, from which an alternative backbone *x*_alt_ is selected. (2) The pair {*x*_ref_, *x*_alt_} is passed to a multi-state inverse-folding model (IF), which generates a set of candidate switch sequences *S*. (3) Each candidate sequence is evaluated with AlphaFold3 (AF3) under both trigger conditions (e.g. with and without phosphorylation) to assess whether the predicted structures recapitulate the two intended conformations.

### 2.2 De novo protein design

De novo protein design aims to create new amino-acid sequences that perform specific functions, such as binding a designated target or catalyzing a specific reaction [Yang et al., 2026]. Modern de novo design pipelines typically proceed in three stages: (1) backbone generation, (2) sequence design and (3) structure-prediction. First, candidate backbone structures capable of supporting the desired function are generated by sampling from a diffusion-based generative model (Section 2.3) trained on large collections of experimentally determined protein structures from the Protein Data Bank (PDB) [Berman et al., 2000]. These backbones provide a structural pool for the second stage, sequence design, in which inverse-folding (IF) models such as ProteinMPNN [Dauparas et al., 2022] and Caliby [Shuai et al., 2025] assign amino-acid sequences expected to fold into each candidate backbone. In the final stage, candidate sequences are ranked or filtered to identify a small subset for experimental validation, which is typically the limiting step. Structure-prediction models such as AlphaFold3 [AF3, Abramson et al., 2024] are widely used for this purpose and have shown strong performance in filtering designs that fail to achieve the desired structural properties.

Our work focuses on the backbone-generation step of this pipeline: in Section 3 we develop a sampling procedure that biases the backbone diffusion model toward ensembles compatible with switch behaviour, leaving the inverse-folding and filtering stages unchanged.

### 2.3 Protein backbone diffusion models and inference-time control

Diffusion models [Ho et al., 2020, Song et al., 2021, Karras et al., 2022] are generative models that learn to sample from a target distribution by progressively denoising Gaussian noise: starting from pure noise, a learned denoising network is iteratively applied across many small steps until a clean sample emerges. Trained on protein backbone structures from the PDB, they form the basis of most generative models used for de novo backbone design [Watson et al., 2023, Lin et al., 2024, Geffner et al., 2025, Yim et al., 2023, Team et al., 2025]. The pretrained backbone diffusion model plays the role of a prior *p*(*x*) over backbones *x* ∈X: it determines what counts as a physically realistic backbone and intuitively defines the feasible set within which our designs must lie.

In design tasks, samples from *p* on their own are rarely useful. We typically have a target property in mind, captured by a reward function *r*: X → ℝ that scores candidate backbones according to the design objective. The natural object to sample from is then the reward-tilted posterior distribution,

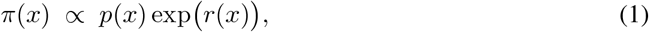

in which the prior continues to provide structural realism while the exponential tilt concentrates mass on configurations that score highly under *r*. Several methods are available to sample from *π* at inference time, using only the pretrained denoiser and the reward [Chung et al., 2022, Dhariwal and Nichol, 2021, Bansal et al., 2023, Wu et al., 2023, Skreta et al., 2025, Singhal et al., 2025, He et al., 2025]; we use the twisted diffusion sampler (TDS) of Wu et al. [2023], which combines reward-guided diffusion with sequential Monte Carlo [Del Moral et al., 2006] over a batch of *N* particles and is exact for *π* in the *N* → ∞ limit. We refer the reader to Appendix A for details.

### 2.4 Well-tempered metadynamics

Metadynamics [Laio and Parrinello, 2002] is an enhanced-sampling technique developed in the molecular dynamics (MD) community to accelerate exploration of complex biomolecular free-energy landscapes [Hénin et al., 2022]. The method projects the system’s dynamics onto a small set of bias potential *V* (*ξ*) on the CV marginal *p*(*ξ*) = *p*(*x*) δ(*ξ* − *ξ*(*x*)) d*x*, repelling the system from collective variables (CVs) *ξ* : → X ℝ^*d*^ (*d* ≪ dim X) and progressively builds a history-dependent bias potential V (ξ) on the CV marginal p(ξ) = R p(x) δ(ξ − ξ(x)) dx, repelling the system from previously sampled regions and promoting transitions between metastable states.

In well-tempered metadynamics [Barducci et al., 2008] the bias is built up in batches. Starting from *V*_0_(*ξ*) = 0, given a batch 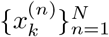 of *N* samples drawn under *V*_*k*_, the bias is updated by depositing one tempered Gaussian hill per sample,

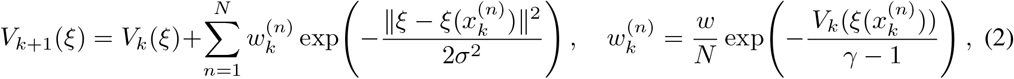

with kernel width *σ*, initial hill height *w*. The bias factor *γ >* 1 controls how aggressively further hills are suppressed in already-sampled regions: low *γ* gives rapid convergence but may under-explore, while high *γ* gives more thorough exploration at the cost of slower convergence. As *k*→ ∞ the marginal of *ξ* under the biased distribution converges to the well-tempered target *p*^wt^(*ξ*) ∝ *p*(*ξ*)^1*/γ*^ [Barducci et al., 2008, *Invernizzi and Parrinello*, *2020]*, *which flattens the basins of ξ* without erasing them. Further details on well-tempered metadynamics are given in Appendix B.

## 3 De novo protein switch design

Despite significant advances in de novo protein design, the design of protein switches from scratch remains a major challenge. This difficulty arises from the stringent requirements imposed on proteins that must reversibly switch between two well-defined, fully folded conformations. Such proteins typically undergo large conformational rearrangements. Often, one conformation can be described as closed, characterised by close contacts between residues in distinct protein domains, while the alternative open conformation involves separation of these domains. In biological systems, transitions between conformations are triggered by effectors, which alter the relative statistical weights with which different conformations are sampled. From a de novo design perspective, this translates into the need to (i) define at least two distinct, interconvertible conformations, (ii) specify a trigger that biases the conformational equilibrium, and (iii) identify a single amino-acid sequence whose energy landscape encodes all of these features.

Conventional de novo protein design pipelines are built on the assumption of a one-backbone-one-sequence relationship [Abrudan et al., 2026], which provides no direct mechanism for the multi-backbone-single-sequence mapping required for switches; existing approaches therefore rely heavily on expert knowledge, manual structural manipulation, and force-field relaxation to construct backbone conformations that are intuitively compatible with a single sequence [Langan et al., 2019, Buckley et al., 2025]. In contrast, protein backbone diffusion models are pre-trained on large-scale structural datasets containing many examples of domain motion and autoinhibitory proteins, suggesting that they may implicitly capture structural features relevant to switch design. This observation motivates our approach: we use a pretrained backbone diffusion model to sample diverse ensembles of closely related backbones that differ primarily in the relative arrangement of individual domains. To support switch design, sampled ensembles must satisfy two criteria: (1) high local structural similarity within each domain, and (2) diversity in domain-level arrangements. Motivated by this, we propose *Diff-Switch*, addressing both requirements in a unified and automatic way. In the following, we will first describe and formulate each requirement, and then combine them in Section 3.3.

### 3.1 Enforcing Local Structural Similarity within Each Domain

To preserve the local structure of each domain, we instantiate a reward function as

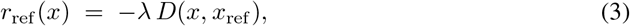

where *x*_ref_∈ X is a reference backbone, *D*(, *x*_ref_) a domain-wise structural distance to *x*_ref_ (for instance, the sum of per-domain RMSDs after rigid alignment of each domain), and *λ >* 0 controls how strictly each domain must match the reference. The corresponding target is

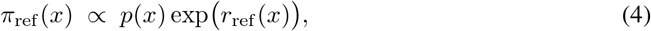

from which we draw a batch of *N* backbones using twisted diffusion sampler (TDS), as described in Section 2.3. As *D* is computed within each domain rather than on the full structure, *π*_ref_ places mass on backbones that share local geometry with *x*_ref_ but may differ arbitrarily in the global arrangement.

### 3.2 Enforcing Global Structural Dissimilarity between Switches

Targeting *π*_ref_ alone does not produce the ensemble diversity we want. The locally-tilted prior is multimodal in the global arrangement of the domains, with modes separated by free-energy barriers that TDS, like any direct sampler, struggles to cross within a finite particle budget; repeated batches therefore concentrate on the dominant global mode of *π*_ref_ and rarely visit the alternative arrangements that are essential for switch behaviour. Compounding this, *p* is determined by the data on which the diffusion model was trained, and physical configurations relevant to a given switch may lie in low-density regions of the prior, so that *π*_ref_ inherits this limited coverage.

To enforce diversity at the global level, we bring the idea of metadynamics as discussed in Section 2.4 into the generation process. Concretely, we add a history-dependent bias on a one-dimensional collective variable *ξ* : X → ℝ (the special case *d* = 1 of the construction in Section 2.4) summarising the global feature along which switch states should differ. The choice of *ξ* is not incidental: it determines the family of switch mechanisms targeted by the resulting ensemble, and natural choices include the hinge angle of a hinge-motion switch and the centre-of-mass distance or relative orientation between two rigid domains. In the context of de novo switch design, the choice of *ξ* ultimately depends on the structural transition one aims to realise. From an energetic perspective, the optimal collective variable should coincide with the true reaction coordinate associated with the slow, functionally relevant motion connecting the conformational states. However, such reaction coordinates are generally not known a priori, and practical choices of *ξ* are therefore guided by structural intuition about the desired conformational change.

### 3.3 Diff-Switch

Diff-Switch iterates the local-similarity tilt of Section 3.1 and the well-tempered metadynamics bias of Section 3.2. Starting from *V*_0_ ≡ 0 so that *π*_0_ = *π*_ref_ recovers the locally-tilted prior, at each iteration *k* = 0, 1, …, *K* − 1 we draw a batch of *N* backbones from the iteration-*k* target

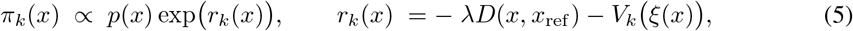

using TDS (Section 2.3), and update the bias from *V*_*k*_ to *V*_*k*+1_ via (2). Inheriting the well-tempered convergence guarantee of Ssection 2.4, the marginal of *ξ* under *π*_*k*_ converges to *π*_ref_ (*ξ*)^1*/γ*^ as *k* → ∞: the sampler progressively traverses the CV-space basins of the locally-tilted prior at a rate set by the bias factor *γ*. This convergence statement applies here because the local-similarity term keeps each domain close to its reference geometry across the support of *π*_*k*_, so *D*(*x, x*_ref_) varies primarily along

#### Algorithm 1

Diff-Switch.

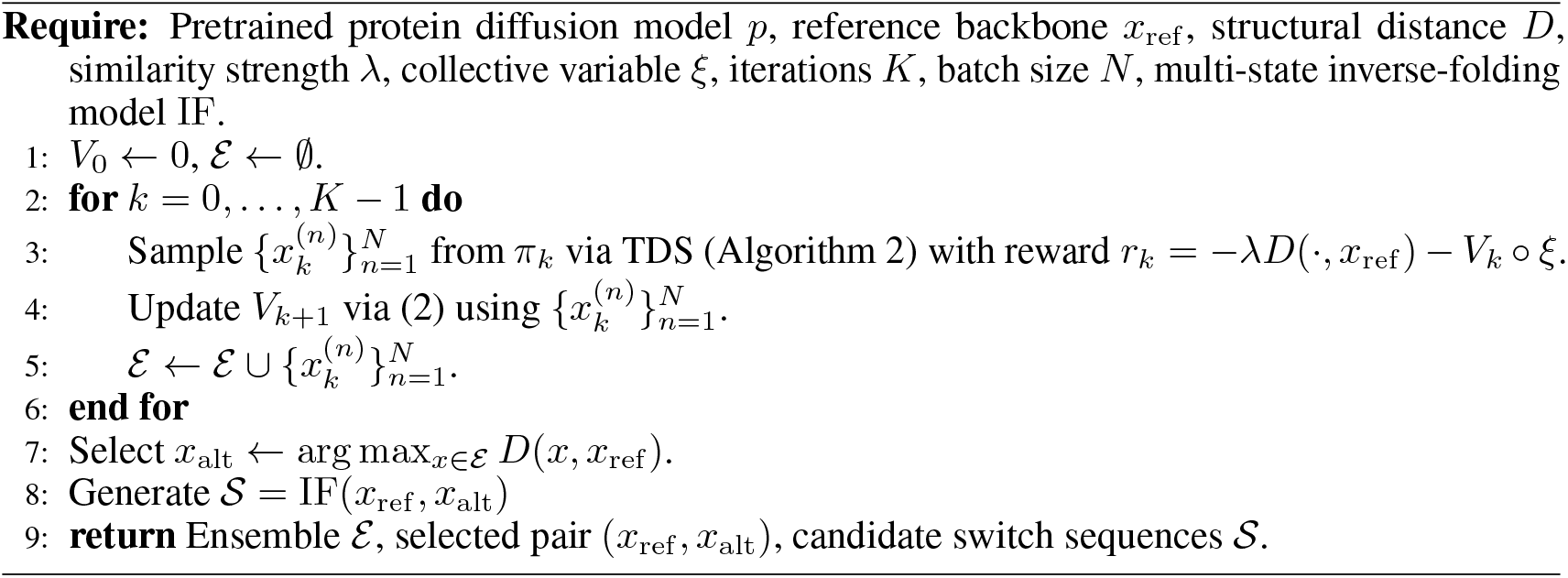

the inter-domain directions captured by *ξ*; in that regime exp(*r*_ref_ (*x*)) acts as a near-constant offset along *ξ*, and *V*_*k*_ can be interpreted directly as a metadynamics bias on the CV-space marginal of *π*_ref_.

After *K* iterations we collect every sampled backbone into an ensemble 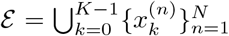 of size *NK*. Multi-state inverse folding requires only a small number of conformations as input, so we select from the alternative backbone *x*_alt_ most dissimilar to *x*_ref_, measured by dissimilarity metric *D*(*x, x*_ref_) (e.g. RMSD or a CV-space distance). The pair *x*_ref_, *x*_alt_ represents the two conformations the designed switch should adopt, and is passed to a multi-state IF model such as ProteinMPNN-MSD [Wicky et al., 2022] or Caliby [Shuai et al., 2025] to obtain an ensemble *S* = IF(*x*_ref_, *x*_alt_) of candidate switch sequences. The sequences in can then be filtered using a structure-prediction model such as AlphaFold3 [Abramson et al., 2024]. The full procedure is summarised in Algorithm 1 and Figure 1.

## 4 Related works

Combining metadynamics with pretrained diffusion models is an actively developing area [Nam et al., 2025, Xie et al., 2026, Lam et al., 2026, Ohnuki and Okazaki, 2025]. Closest to and concurrent with our work, Xie et al. [2026] use well-tempered metadynamics with a pretrained diffusion model for inference-time control, targeting rare-event sampling and free-energy estimation in molecular systems rather than protein switch design.

Existing de novo switch designs have used relatively simple strategies across the three stages of protein design. To engineer a hinge-based conformational switch, Praetorius et al. [2023] started from a tightly packed helical repeat protein (DHR) backbone [Brunette et al., 2015] representing the closed state, and manually generated an open conformation through a sequence of shifting, rotation, stitching, and Rosetta-based relaxation [Leaver-Fay et al., 2011]. Both conformations were then provided to ProteinMPNN multi-state design [Wicky et al., 2022] for sequence design, and switching behaviour was evaluated using AlphaFold2 with initial guess (AF-IG) [Bennett et al., 2023], run with and without the effector peptide.

To design phosphorylation-induced protein switches, Buckley et al. [2025] used compact de novo protein folds in which a phosphorylation site is buried in the closed state. Rather than constructing an open conformation explicitly, they masked the helical terminus containing the phosphorylation site, made possible by their use of a prior amino-acid sequence for that terminus suitable for downstream experiments. ProteinMPNN-MSD [Wicky et al., 2022] was used for sequence optimization, and AF3 was used to assess whether phosphorylation stabilized the open state. The general design purpose of this work motivated our experiments in Section 5.2.

## 5 Experiments

In this section, we evaluate the utility of our generated backbone ensembles for designing sequences that satisfy the structural diversity of a protein switch.

### 5.1 Validation on domain-rearrangement protein switches

We make use of a seminal dataset of two-state domain-motion proteins, simulating the motion of domain-rearrangement protein switches [Kalakoti and Wallner, 2025]. This is a diverse dataset that consists of 20 proteins from 109 to 801 amino acids long (average: 398) with distinct open and closed conformations, both of which are available in the Protein Data Bank (PDB), and contains various switch mechanisms from ligand binding to enzymatic active site closure. To simulate a real-world protein switch design scenario where only one state is known, we provide our method with a single starting backbone conformation, i.e. the backbone structure of either the closed or the open conformation. Our method then samples a diverse ensemble of backbone structures, effectively containing the starting conformation (closed or open) and conformations of the alternative state (open or closed) as well as the transitional states between open and closed ones. This is done by first selecting a set of CVs capable of describing the coordinates along which switch reaction occurs. We choose a one-dimensional CV, center-of-mass distance between each protein’s domains, which intuitively is capable of separating the open and closed conformations.

For each protein we compare two design strategies. In the baseline, the reference backbone *x*_ref_ is fed directly to the inverse-folding method to generate candidate sequences. Even with a single backbone, this can occasionally yield switch-compatible sequences, because proteins are inherently dynamic and a sequence optimized for one reference can transiently access conformational states resembling alternative functional geometries. Diff-Switch instead identifies an alternative backbone *x*_alt_ from the sampled ensemble, which is paired with *x*_ref_ as input to the inverse-folding method, producing an ensemble of candidate sequences *S*. We then use AF3 to generate the predicted structure. See Figure 2 for an illustrating example of the backbones generated by Diff-Switch.

**Figure 2.**
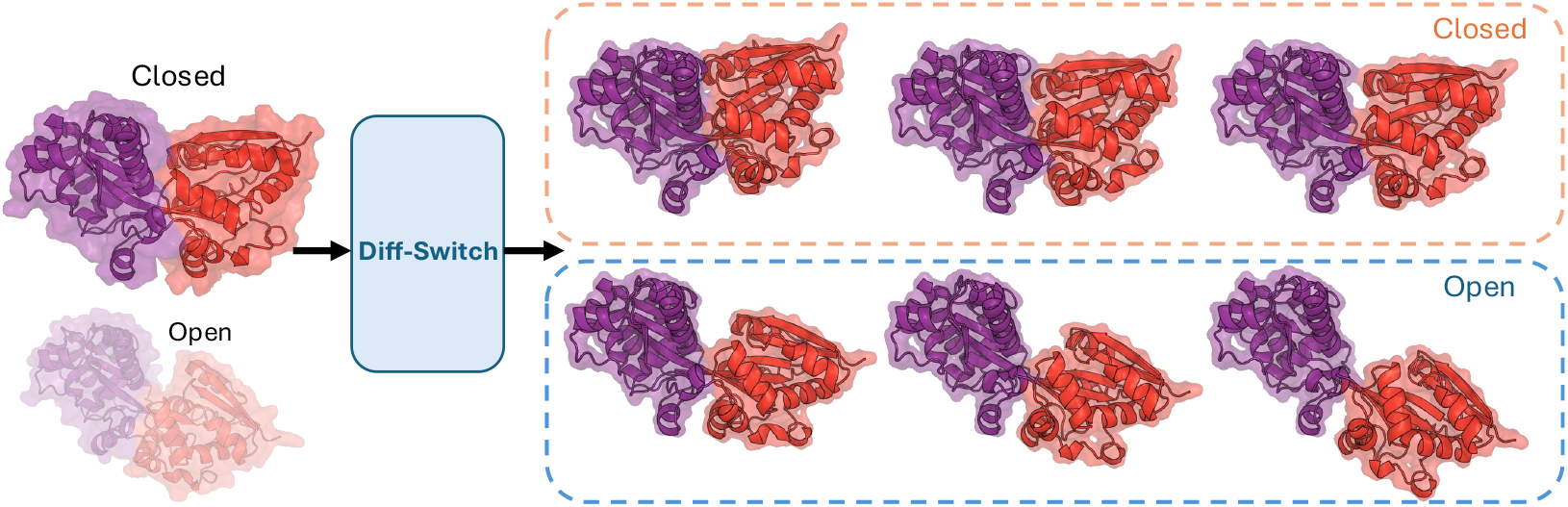
Conformational diversity generated by Diff-Switch for one system in the dataset (P31133). The closed conformation (top left) is provided to the pipeline as the reference *x*_ref_, with the open ground-truth conformation (bottom left) shown for comparison. Six examples of backbone structures emerged from the pipeline (right) recover both target conformations despite only the closed reference being provided as input. The bottom-right backbone structure (the most-open one) is identified as *x*_alt_ since it is the most-different structure to the *x*_ref_. Structures are shown in ribbon (solid) and surface (transparent) representations; the two enforced local-structure domains are coloured in red and purple.

We evaluate whether a designed sequence *s* ∈ *S* supports both switch endpoints using self-consistency metric, scRMSD_switch_ and scTM_switch_. Following the evaluation pipeline of Abrudan et al. [2026], we run AlphaFold3 separately on *s* together with each of the open and closed ground-truth backbones, and compute self-consistency between each predicted structure and the corresponding ground truth, i.e. scRMSD_open_ and scRMSD_closed_ as well as scTM_open_ and scTM_closed_. scRMSD_open_ and scTM_open_ show if *s* has the capacity of folding into the open ground-truth backbone. Similarly, scRMSD_closed_ and scTM_closed_ indicate compatibility of sequence *s* with the closed ground-truth backbone. Because a switch sequence must fold into both backbones, we take the worse of the two values (the higher scRMSD and the lower scTM) as the performance metric for *s*. Full details are given in Appendix D.

Diff-Switch outperforms the baseline in every configuration we test (Figure 3). The area under the success-rate curve increases by between 0.09 and 0.17 across the four combinations of reference conformation (open, closed) and inverse-folding backend (ProteinMPNN, Caliby), with the largest gain (0.51→ 0.68, in the closed-reference Caliby setting) closing most of the Caliby-ProteinMPNN gap. We used a single set of Diff-Switch hyperparameters across all 20 proteins and both inverse-folding backends (Appendix C); the per-protein variability visible on the right of Figure 3 therefore reflects the inherent volatility of the base diffusion model rather than tuning differences. One-sided paired Wilcoxon signed-rank tests over the 20 proteins confirmed that Diff-Switch significantly improves both metrics relative to the single-conformation baseline in every setting: for Caliby, *p* = 0.002*/*0.005 for closed/open scRMSD_switch_ and *p* = 0.003*/*0.012 for closed/open scTM_switch_; for ProteinMPNN, *p* = 0.002*/*0.016 for closed/open scRMSD_switch_ and *p* = 0.004*/*0.027 for closed/open scTM_switch_. We view the numbers in this evaluation as a floor rather than a ceiling: per-problem tuning of *ξ, λ*, and the metadynamics schedule, together with stronger pretrained backbone priors and inverse-folding models, should push performance further.

**Figure 3.**
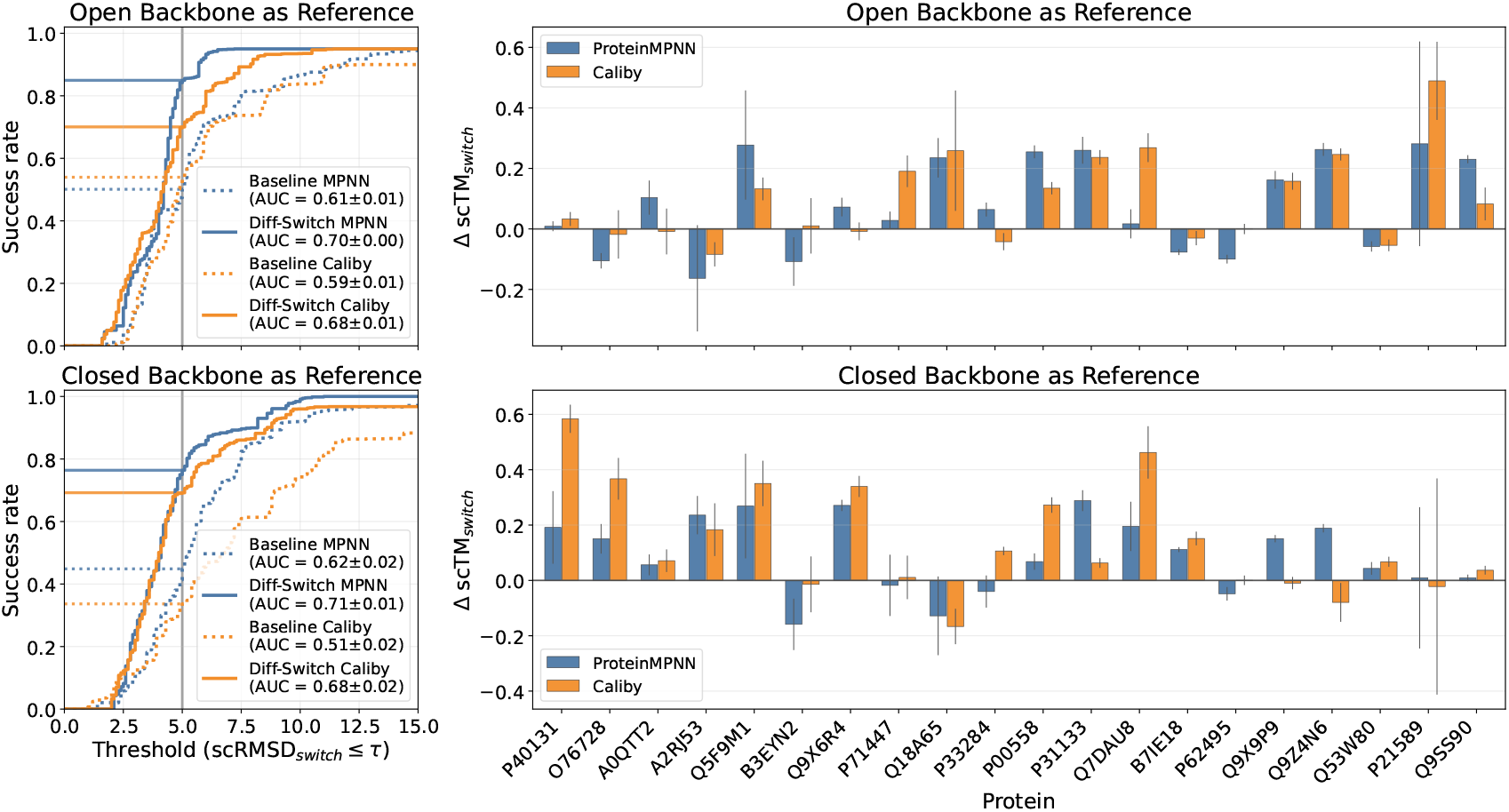
Evaluation of the performance of Diff-Switch on domain-rearrangement protein switches. **Left:** Success rates defined by different scRMSD_switch_ thresholds *τ* for predicted structures of baseline sequences (dotted) and those generated for backbones emerging from Diff-Switch (solid). Sequences were generated using either ProteinMPNN (blue) or Caliby (orange). The vertical line marks the threshold *τ* = 5 Å (see Appendix D) below which a predicted structure is considered to match the target conformation. Bootstrapping is done (see Appendix D) on predicted structures and AUCs are reported in the legend. **Right:** Differences between scTM_switch_ scores calculated for structures predicted from sequences that were generated using the Diff-Switch pipeline and the baseline sequences. Positive values indicate better agreement with experimental structures for sequences emerging from the Diff-Switch pipeline. Error bars indicate the standard deviation of bootstraps. We investigate the noisy case P21589 in Figure 5.

### 5.2 De novo design of phosphorylation-induced protein switches

We next evaluate Diff-Switch on a more targeted switch-design setting: phosphorylation-induced conformational switching. In this class of systems, a protein is designed so that phosphorylation of a designated residue destabilizes one conformation and shifts the conformational equilibrium toward an alternative state. Following the design logic of Buckley et al. [2025], we focus on compact protein backbones in which the phosphorylation site is buried in the closed state. Upon phosphorylation, the added negative charge is expected to favor an open conformation in which the modified residue becomes solvent exposed and avoids unfavorable local packing interactions.

In this task, only the closed conformation is available and the open conformation must be generated rather than taken from an experimentally resolved conformational pair. Also, the desired switching coordinate is tied to a chemically interpretable trigger, namely exposure and displacement of the phosphorylation-site-containing segment. We therefore use Diff-Switch to sample alternative backbones that preserve the local geometry of the folded domains while encouraging motion of the phosphorylation-site region away from the protein core. In practice, we define the collective variable as the center-of-mass distance between the phosphorylation-site helix and the remaining scaffold. The resulting ensemble is filtered to select a candidate open conformation that maintains high local structural quality while increasing exposure of the phosphorylation site.

For a selected closed/open backbone pair, we run Caliby to design 16 sequences compatible with both conformations. We then evaluate candidate sequences using structure prediction in the unmodified and phosphorylated settings. A successful design should satisfy two criteria: in the absence of phosphorylation, the predicted structure should remain close to the intended closed state, keeping the interface between the scaffold and the helix packed; whereas in the phosphorylated setting, the predicted structure should open up and completely expose the interface, available for potential molecule partners to bind it (Figure 4). To quantify the criteria, we report two complementary metrics under both with and without phosphorylation conditions for each designed sequence: the center-of-mass distance between the phosphorylation-site helix and the scaffold, and the buried interface area between them. The former measures how open a structure is and the latter expresses the accessibility of the interface before and after the phosphorylation. Looking at Figure 4, we seek designed sequences that have low center-of-mass distances (high buried interface areas) without phosphorylation, and high center-of-mass distances (near zero buried interface areas) with phosphorylation. 25% (4*/*16) of the Diff-Switch designed sequences showed switch behavior while none of the 16 sequences we designed using only the closed conformation as input to Caliby exhibited this behavior. Figure 1 summarizes this pipeline and representative designs.

**Figure 4.**
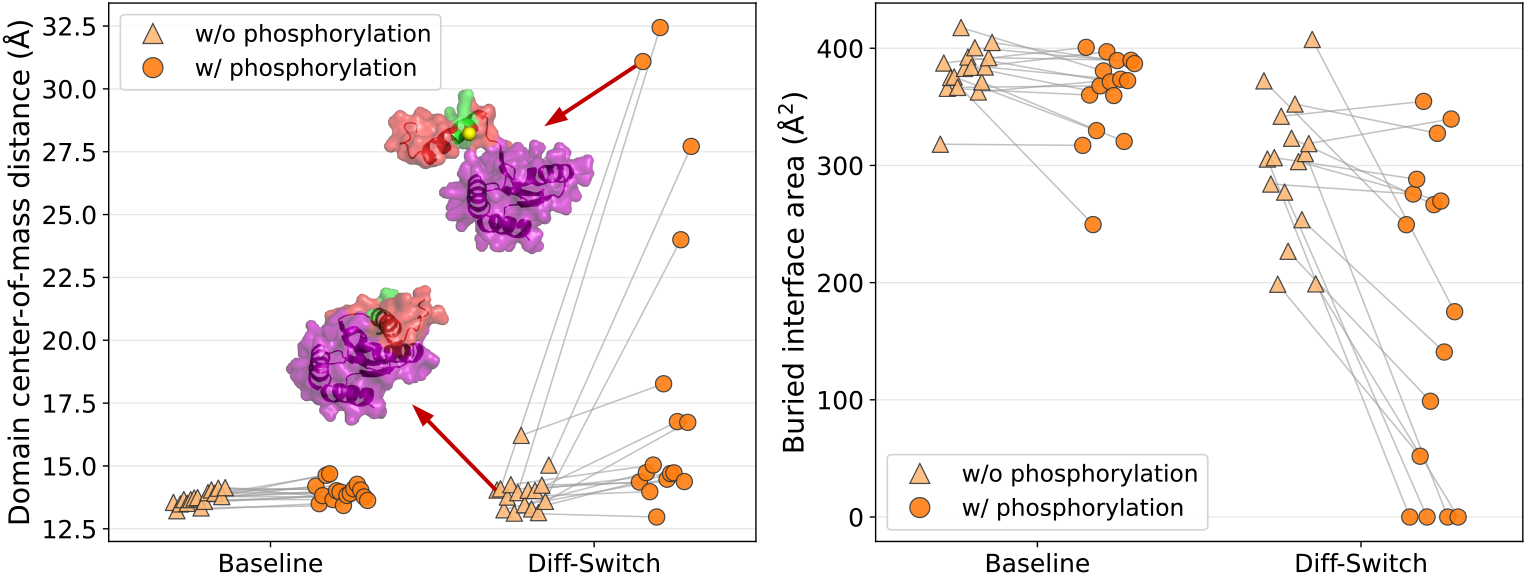
Validation of phosphorylation-induced protein switches from Diff-Switch. **Left:** Domain center-of-mass distance (Å) between the phosphorylation-site helix (residues 90-102) and the remaining scaffold (residues 1-83), shown per designed sequence under two AF3 conditions: w/o phosphorylation (unmodified; light orange triangles) and w/ phosphorylation (Residue 95 converted from serine to phosphoserine; dark orange circles). Points from the same sequence are connected by thin gray lines. Sequences designed with Diff-Switch are compared with Baseline on the horizontal axis. The structures of a successful switch design are overlaid on the plot; the scaffold is colored in purple and the helix is in red and green. **Right:** Same layout, but showing the buried interface area (Å^2^) of the scaffold interface residues, i.e. the scaffold surface occluded by the phosphorylation-site helix, computed as the change in solvent-accessible surface area (SASA). Larger values indicate a more closed interface; zero means the interface is completely exposed.

Unlike Section 5.1, this setting is more realistic: only the closed conformation is provided, and the open state must be discovered rather than taken from a pre-existing experimental pair. Diff-Switch removes the manual structural intervention used by Buckley et al. [2025] (where the phosphorylation-site terminus is masked to construct the open state) and replaces it with an automatic CV-driven search, demonstrating that the method generalises to settings where no experimental conformational pair is available.

## 6 Discussion

We introduce Diff-Switch as a principled alternative to existing protein switch design approaches by reframing switch design as controlled ensemble generation. Our method uses metadynamics to impose a history-dependent bias along a user-defined switch coordinate, enabling direct control over the structural dimension along which conformations diversify. This provides a unified and scalable framework for targeting a wide range of switching behaviors, from hinge motions to more complex domain rearrangements. Moreover, the same metadynamics formulation can be used to estimate relative free-energy differences along the chosen coordinate, linking generative modeling with energetic characterization.

While metadynamics has long been used to capture slow, functional motions, the corresponding reaction coordinates are typically not known a priori. In our test cases (Section 5.1), a simple geometric choice, the center-of-mass distance between domains, proved effective in the majority of cases, despite being a crude proxy for the true reaction coordinate. It enabled the generation of backbone ensembles that supported the design of sequences capable of more closely adopting both open and closed conformations. Notably, this behavior was observed when initializing from either the open or the closed state, highlighting the robustness of the approach.

Our choice of collective variable, the centre-of-mass distance between domains, was selected to align with the typical common open/close switching motion observed in proteins. In a small number of proteins this choice fails: the underlying transition involves inter-domain rotations that leave the centre-of-mass distance approximately unchanged, and the metadynamics bias therefore cannot drive the sampler toward the alternative state. Figure 5 shows one such case. Even on the affected proteins, however, the impact is contained: Diff-Switch falls below baseline on scTM_switch_ only in the closed-reference, Caliby configuration.

**Figure 5.**
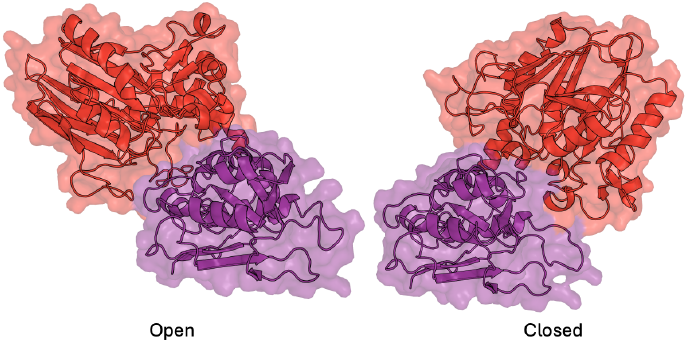
Open and closed conformations of P21589, an example where the centre-of-mass distance between domains fails as a collective variable. The two states differ primarily by an inter-domain rotation that leaves the centre-of-mass distance approximately unchanged, so the metadynamics bias on *ξ* cannot drive the sampler across the conformational transition. This leads to unstable performance in Figure 3 (Right).

It is important to emphasize that, in a de novo design setting, the choice of CV must reflect the desired switching behavior specified by the designer. For example, if we desire a protein domain to rotate instead of opening, the center-of-mass distance is a sub-optimal choice. However, in Section 5.2, Diff-Switch with the center-of-mass distance CV enables the generation of backbone conformations consistent with the expected structural effect of phosphorylation, namely altered interactions between a modified helix and the rest of the protein. Accordingly, the collective variable was chosen to reflect this anticipated structural change. While similar triggers have been used in prior work, they have typically been incorporated in a more heuristic manner.

## 7 Conclusion

In this work, we introduce Diff-Switch, a framework that reframes protein switch design as controlled ensemble generation. By combining metadynamics with pretrained diffusion models, it enables scalable, inference-time control of conformational diversity. Coupled with multi-state inverse folding, Diff-Switch yields sequences whose predicted conformations get close to experimentally observed switch states. More broadly, it highlights how controllable generative models can bridge structure, dynamics, and function in protein design.

### Limitations and future work

A key limiting factor of our method is the choice of the CV, which must capture and constrain the intended structural transition. Identifying appropriate CV remains an active area of research across many domains, and advances in this area will directly benefit Diff-Switch. In its current formulation, the method primarily supports rearrangements of rigid or semi-rigid domains. More complex transitions, such as fold changes involving the formation or loss of secondary structure (e.g., helix–sheet transitions or disorder–order transitions), are not captured. Finally, a scalable de novo protein switch design pipeline also requires systematic approaches for designing the trigger or stimulus that induces the conformational change. While we demonstrate this for phosphorylation in Section 5.2, extending the approach to other triggers, such as protein–protein interactions, remains an important direction for future work.

## Acknowledgments and Disclosure of Funding

JH acknowledges support from the University of Cambridge Harding Distinguished Postgraduate Scholars Programme. SS acknowledges the support of a CANSSI CRT grant and the NSERC Discovery Grant.

## A Protein backbone diffusion and Twisted Diffusion Sampler

This appendix describes the twisted diffusion sampler (TDS) of Wu et al. [2023] as we instantiate it for protein backbone design. We first sketch how diffusion models are typically constructed for protein backbones, then specify the discrete-time noising and denoising chain that defines the prior *p*, develop the annealed sequence of intermediate targets, the reward-guided proposal, and the SMC importance weights that constitute TDS, and close with a comparison to Feynman–Kac-style correctors that motivates our choice.

### A.1 Diffusion models for protein backbones

The prior *p* used throughout the paper is a pretrained generative diffusion model on the backbone space X = ℝ^*L×*4*×*3^ defined in Section 2.2. Modern backbone diffusion models [Lin et al., 2024, Geffner et al., 2025, Yim et al., 2023, Team et al., 2025] are constructed to be invariant under global SE(3) transformations of the backbone, either by using SE(3)-equivariant networks on Cartesian coordinates or by operating on per-residue rigid frames in SE(3)^*L*^. For the constructions in this appendix the architectural choice is immaterial: we treat the prior as a generic denoiser-based diffusion model, and the TDS algorithm below depends only on the denoiser, applying whenever the prior is presented in this form.

### A.2 Discrete-time noising and denoising chain

A clean backbone *x*_0_ ~ *p* is perturbed by Gaussian noise to give *x*_*t*_ = *x*_0_ + *tϵ* with *ϵ* ~ *N* (0, *I*), yielding marginals *p*_*t*_ that interpolate between *p* at *t* = 0 and approximately *N* (0, *T* ^2^*I*) at *t* = *T*. We denote the conditional mean by 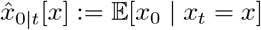, which can be approximated by a neural network learned from samples; throughout this appendix we treat 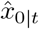 as given. It is related to the score by Tweedie’s identity, 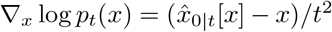. Sampling proceeds along a discretized noise schedule

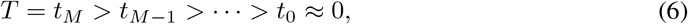

which induces a forward Markov chain with Gaussian transition kernel

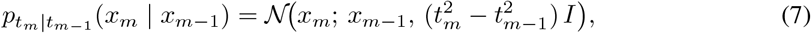

and a discrete-time reverse-kernel approximation parametrized by 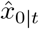,

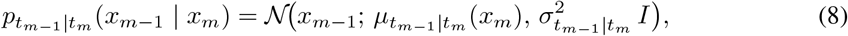

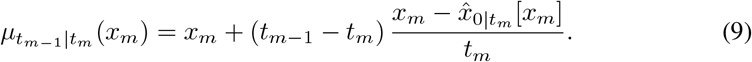

The per-step standard deviation 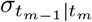 is a hyperparameter of the sampler: setting 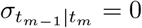 yields a deterministic ODE-flow update, while non-zero values give stochastic samplers such as the EDM scheme of Karras et al. [2022]. Iterating (8) from 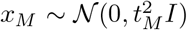 down to *x*_0_ defines the discrete-time distribution induced by the pretrained sampler, which we identify with the prior *p*. The TDS construction below extends to other denoiser-based parametrisations with appropriate modifications to the kernel forms.

### A.3 Annealed sequence of intermediate targets

Given a reward *r* : X → ℝ, our objective is the reward-tilted target *π*(*x*) ∝ *p*(*x*) exp(*r*(*x*)) from (1). Following the TDS construction [Wu et al., 2023], we define a sequence of intermediate targets indexed by the noise schedule,

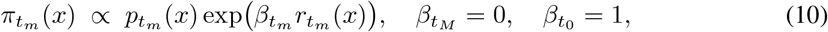

Where 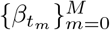 is a non-decreasing annealing schedule and

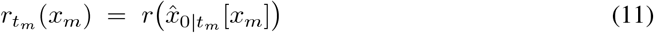

is a noise-level twist that evaluates the reward on the predicted clean structure 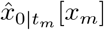. By construction, 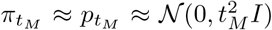 is easy to sample from and 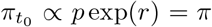 recovers the desired tilted target. Annealing the reward in this way avoids the variance collapse that arises from applying the full reward at the noisiest end of the chain, where the predicted clean structure is uninformative.

### A.4 Reward-guided proposal kernel

To bias the reverse-time chain toward high-reward regions, we replace 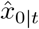 with a reward-guided prediction 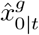 in the reverse-kernel update, yielding a proposal kernel to guide the particle towards regions of higher reward,

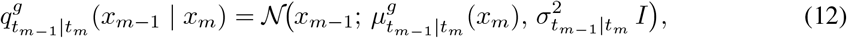

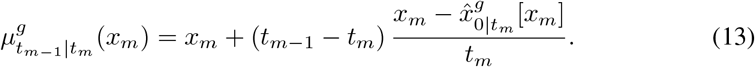

A standard choice is the additive gradient correction

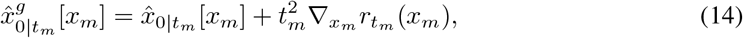

which corresponds to guidance for diffusion posterior sampling [Dhariwal and Nichol, 2021, Chung et al., 2022]. The TDS construction is, however, agnostic to the precise form of 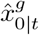 : any guided prediction that concentrates probability near high-reward regions can be used as a proposal, and the importance weights below correct for the resulting bias.

### A.5 SMC importance weights and resampling

Because the guided proposal in (12) does not generally coincide with the kernel that targets 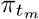, we correct the discrepancy with sequential Monte Carlo. At each transition *x*_*m*_ → *x*_*m*−1_, the incremental log-weight is

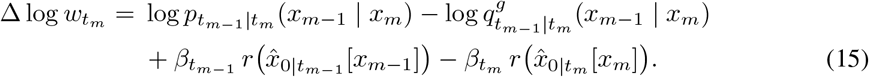

The first line is the standard importance ratio between the base reverse kernel and the guided proposal; the second line is a telescoping reward increment that anneals from 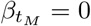 at the noisiest end of the chain to 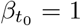 at the clean end. With *N* particles 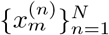 propagated independently under (12) and weights updated according to (15), the normalized weights

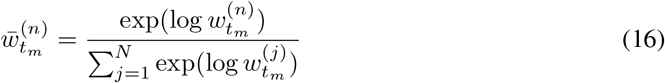

provide a particle approximation to 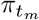 that is asymptotically exact in *N*. To control weight degeneracy we monitor the effective sample size

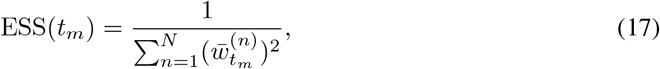

and apply systematic resampling whenever 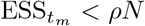 for a user-specified threshold *ρ*∈ (0, 1], resetting log-weights to zero after resampling. The full procedure is summarized in Algorithm 2.

### A.6 Discrete-time exactness and comparison to Feynman-Kac correctors

Several closely related particle-based correction schemes have appeared in the recent literature [Wu et al., 2023, Skreta et al., 2025, Singhal et al., 2025, He et al., 2025]. An alternative to TDS is the Feynman–Kac corrector (FKC) [Skreta et al., 2025], which derives importance weights from a continuous-time Feynman–Kac representation of the reverse SDE. As emphasized in the recent RNE analysis [He et al., 2025], FKC weights are exact only in the idealized regime where the diffusion model is the exact time-reversal of the noising SDE *and* the discretisation step vanishes. In contrast, the TDS weights in (15) are exact with respect to the discrete-time distribution actually induced by the chosen sampler and the specified twist function, regardless of how coarse the noise schedule is or how imperfectly 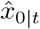 approximates the true conditional mean. Because protein backbone diffusion models are typically run with a small number of denoising steps and finite model capacity, this distinction is practically relevant, and is the main reason we adopt TDS rather than FKC in our pipeline.

#### Algorithm 2

Twisted diffusion sampler.

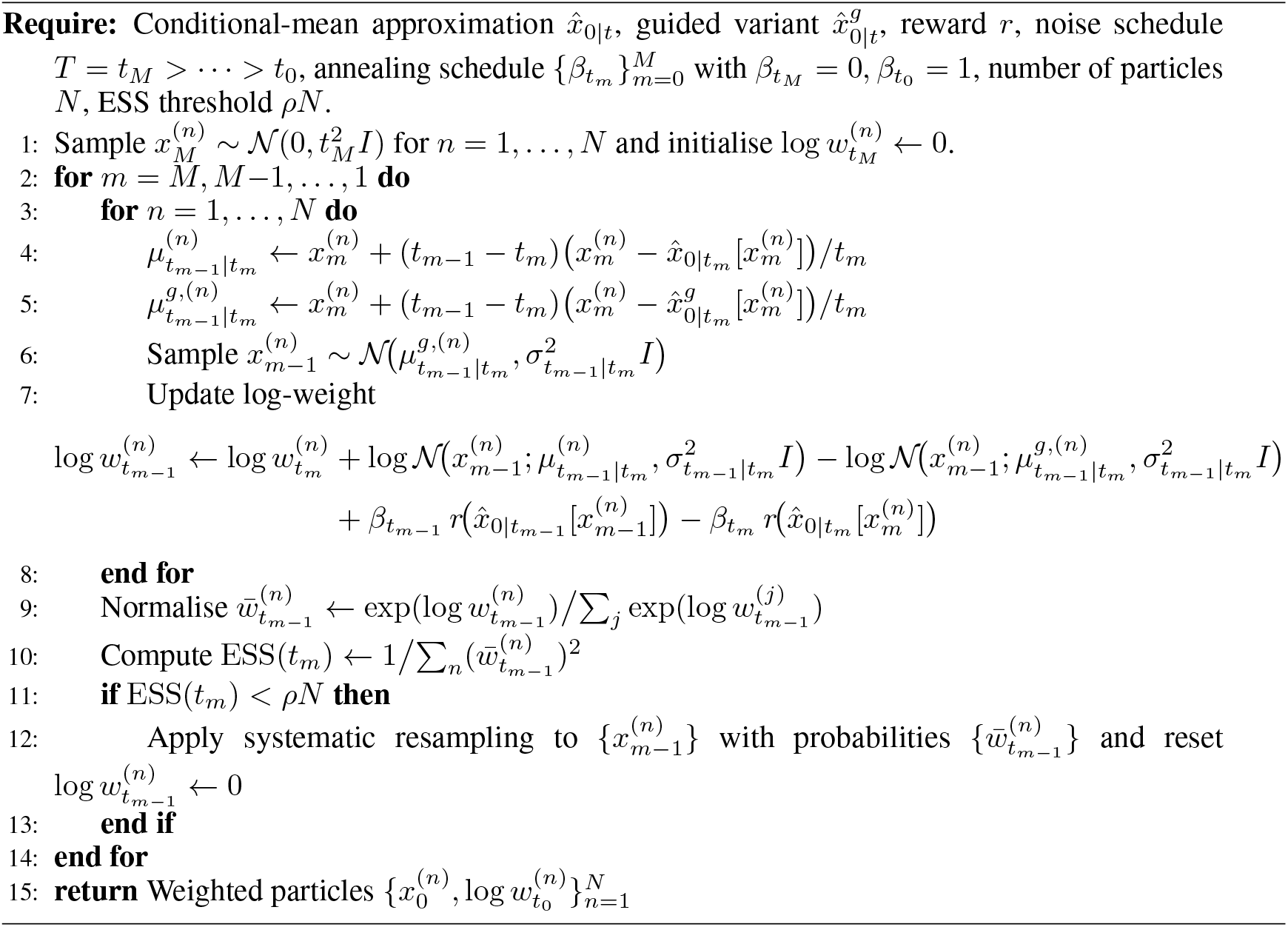

## B Well-tempered metadynamics

This appendix gives a self-contained account of well-tempered metadynamics [Barducci et al., 2008], the variant of metadynamics on which our collective-variable bias is based. We follow the modern presentation of Invernizzi and Parrinello [2020].

### B.1 Collective variables and target distribution

Let *p*(*x*) denote a Boltzmann distribution over molecular configurations *x* ∈ X. A set of *collective variables* is a smooth mapping

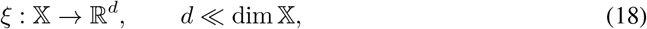

whose components capture the slow degrees of freedom relevant to the conformational changes of interest, such as a hinge angle, a domain centre-of-mass distance, or a torsion angle. The marginal distribution of the CVs under *p* is

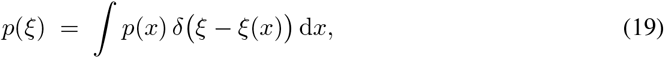

and the associated free energy is defined as *F* (*ξ*) = − log *p*(*ξ*). In MD applications, *p*(*ξ*) is typically multimodal and separated by high free-energy barriers, so direct simulation rarely visits all relevant metastable basins within practical time scales. Metadynamics addresses this by introducing a bias potential *V* (*ξ*) that reshapes the effective marginal toward an easier target 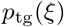, while preserving the conditional distribution *p*(*x* | *ξ*) within each CV slice.

### B.2 Well-tempered target

A natural choice of 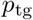 is the well-tempered distribution

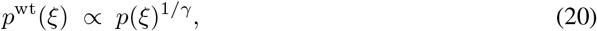

where *γ >* 1 is the *bias factor*. The well-tempered target retains the modal structure of *p*(*ξ*) but flattens the relative populations of metastable basins, making transitions between them more frequent without erasing physically meaningful structure. Two limiting cases give intuition: *γ* → 1 recovers the unbiased dynamics, while *γ* → ∞ corresponds to uniform sampling of CV space.

### B.3 Adaptive hill deposition

Well-tempered metadynamics constructs the bias potential adaptively along the simulation trajectory. Starting from *V*_0_(*ξ*) = 0, at the *k*-th deposition step a sample *x*_*k*_ is drawn under the current bias *V*_*k*_ and a Gaussian hill is deposited at its CV position *ξ*(*x*_*k*_), yielding the updated bias

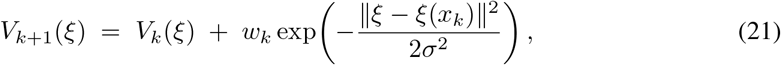

where *σ >* 0 is a fixed kernel width that sets the resolution of the bias in CV space. Unrolling the recursion gives the equivalent expression

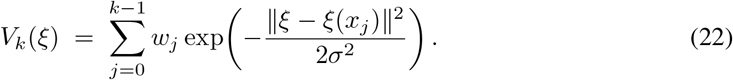

The hill height *w*_*k*_ is itself adaptive, following the well-tempered tempering rule

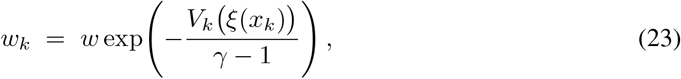

with initial height *w >* 0. The intuition is straightforward: in regions of CV space that have already accumulated substantial bias, *V*_*k*_(*ξ*(*x*_*k*_)) is large and subsequent deposits are exponentially suppressed; in unexplored regions, *V*_*k*_ is small and full-height hills are deposited. As *k*→ ∞, the bias potential converges, in the well-tempered sense, to

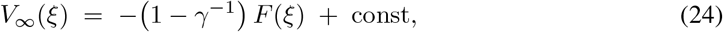

so that the biased marginal distribution coincides with the well-tempered target (20) and the underlying free-energy surface can be recovered, up to an additive constant, from the deposited bias [Barducci et al., 2008].

### B.4 Role of the bias factor

The bias factor *γ* governs the trade-off between exploration and convergence speed. Small values of *γ* lead to aggressive suppression of hills and rapid stabilisation of *V*_*k*_, but can result in under-exploration when the total deposited bias is insufficient to overcome the largest free-energy barriers in CV space. Large values of *γ* allow persistent hill deposition and more thorough exploration, at the cost of slower convergence and noisier free-energy estimates. In the limit *γ*→ ∞ the tempering term in (23) vanishes and one recovers standard (non-tempered) metadynamics, in which hills of constant height *w* are deposited indefinitely. In practice, *γ* is chosen so that *γ* − 1 is comparable to the largest free-energy barrier (in the units of *F*) one expects to traverse in CV space [Barducci et al., 2008].

### B.5 Batched hill deposition

When the dynamics is parallelized across *N* replicas, or, as in our setting, across a population of *N* diffusion particles sharing a common bias, it is convenient to deposit *N* Gaussian hills per update step, one per particle. Given a batch 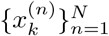 of *N* samples drawn under *V*_*k*_, the bias update becomes

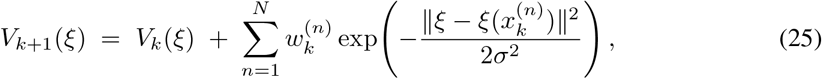

with per-particle tempered hill heights

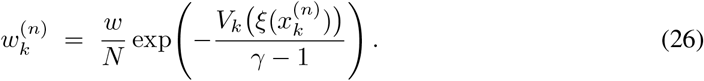

Dividing the initial hill height by *N* keeps the total mass deposited per update step comparable to the single-walker scheme, so that the well-tempered convergence guarantees of Barducci et al. [2008] carry over to the batched setting. This is the form of the bias used in our protein switch sampler.

## C Tuning Diff-Switch

Diff-Switch inherits hyperparameters from three components: the local-similarity reward (Section 3.1), the well-tempered metadynamics bias (Section 3.2 and Appendix B), and the inner TDS sampler (Appendix A). This appendix groups them by component and discusses how each was set in our experiments. Defaults used throughout our experiments are summarized below.

### C.1 Local-similarity reward: *λ* and *D*

The strength *λ >* 0 in *r*_ref_ (*x*) = − *λD*(*x, x*_ref_) controls how tightly the sampler is constrained to backbones that match the reference geometry within each domain. Small *λ* leaves the prior largely unmodified and allows substantial fold variation at the cost of losing the per-domain anchoring required for a single sequence to fold into both states. Large *λ* keeps each domain rigidly close to the reference but suppresses the global rearrangements we want to discover. A useful diagnostic is the per-domain RMSD distribution under *π*_ref_ : *λ* should be set so that this distribution sits comfortably below the typical fluctuation within a folded domain (a few Ångstroms) without becoming so peaked that the sampler explores no global variation. We use *λ* = 3.

The structural distance *D* should evaluate per-domain rather than on the full structure, so that backbones that are locally faithful to the reference but globally rearranged are not penalized. We use the sum of per-domain RMSDs on the backbone atoms of N,C_*α*_,C,and O after rigid alignment of each domain. We did the domain assignments for the benchmark.

### C.2 Collective variable *ξ*

The collective variable is the most consequential design choice in Diff-Switch (Section 3.2). A workable *ξ* must (i) be smooth, differentiable, and cheap to evaluate from a backbone, (ii) resolve the directions along which the two target conformations differ, and (iii) be one-dimensional so that the metadynamics bias can be deposited efficiently with a small number of hills. Our default is the centre-of-mass distance between two pre-defined domains, which works for most domain-rearrangement switches in our experiments. Alternatives we have used or considered include the hinge angle of two-domain hinge motions and a relative-orientation angle between rigid bodies. The phosphorylation case study in Section 5.2 illustrates a setting where the centre-of-mass choice fails and a different *ξ* is required.

### C.3 Metadynamics bias: *σ, w, γ*

Three parameters govern the well-tempered bias deposited on *ξ* (Appendix B). The kernel width *σ* sets the resolution of the bias in CV space and should be small enough to distinguish metastable basins along *ξ* but large enough that successive hills overlap meaningfully; a common heuristic is to take *σ* on the order of one quarter of the typical fluctuation of *ξ* within a basin. The initial hill height *w* and the bias factor *γ* together determine the effective free-energy barrier the bias can flatten: small *γ* favours rapid convergence but risks under-exploration when the largest barrier in CV space exceeds (*γ* − 1) in the units of *F*, while large *γ* allows more thorough traversal of CV space at the cost of slower convergence. In our experiments we use *σ* = 0.3, *w* = 0.1, *γ* = 10.

### C.4 Outer-loop budget: *K* **and** *N*

The number of iterations *K* and the batch size *N* jointly set the compute budget through the total *NK* TDS calls. For fixed *NK*, larger *K* and smaller *N* increase the rate at which the bias *V*_*k*_ is updated, accelerating CV-space exploration but giving each iteration’s ensemble fewer particles to characterise the current target *π*_*k*_. Larger *N* and smaller *K* trade exploration speed for per-iteration sample quality. Small *N* also degrades the asymptotic correctness of TDS as a sampler from *π*_*k*_, since the SMC particle approximation is only exact in the *N* → ∞ limit and the empirical distribution at small *N* remains biased toward the guided proposal rather than *π*_*k*_. For our purposes this is not a binding concern: the guided proposal in TDS already concentrates mass near high-reward regions, so even when the importance weights fail to fully correct for the proposal bias the resulting backbones tend to score well on the reward, which is what design ultimately requires. We use *K* = 20 and *N* = 5, selected so that the deposited bias has approximately stabilized on a held-out reference protein by the final iteration.

### C.5 Inner TDS sampler

The inner sampler inherits the noise schedule *T* = *t*_*M*_ *>* … *> t*_0_ and the number of denoising steps *M* from the pretrained backbone diffusion model: we use the default schedule of [Team et al., 2025, Abramson et al., 2024] with *M* = 400. The reverse-step noise level 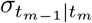 likewise follows the pretrained sampler’s recommendation; we did not retune it. The annealing schedule 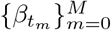 interpolating from 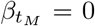 to 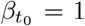 is taken geometric in *m*, which we found sufficient. The ESS resampling threshold is set to *ρ* = 0.5, a standard choice in SMC. The reward-guided prediction 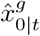 is taken to be the additive gradient correction of (14); we did not experiment with alternative guidance schemes.

### C.6 Selection distance

The distance used in the selection step *x*^*∗*^ = arg max_*x*∈ ℰ_ *D* (*x, x*_ref_) can be either the per-domain RMSD used in the local-similarity reward or the one-dimensional CV-space distance |*ξ*(*x*) − *ξ*(*x*_ref_) |. The CV-space variant is cheaper and aligns the selection with the direction along which we have explicitly biased the sampler; we use it as our default. The per-domain RMSD variant is preferable when the CV is a coarse summary that may pool together backbones that differ along orthogonal directions, in which case picking the most CV-distant backbone need not pick the most structurally distant one.

## D Experimental details

### D.1 Data set

OC23 is a dataset of 23 proteins with distinct open and close structures (TM-Score < 0.85) available in the Protein Data Bank [Kalakoti and Wallner, 2025]. Originally curated to assess the performance of AlphaFold2-based alternative conformation sampling methods, it is well-used in various studies to evaluate multi-state protein prediction and/or design [Richman et al., 2026, Xie et al., 2026, Abrudan et al., 2026]. Out of all the proteins in this set, we dropped A0A075Q0W3 because it is a three-domain protein with no clear closed conformation, and Q9ERE7 due to technical issues with running experiments on it. We also took A6UVT1 out because it is a protein with two open conformations and no closed conformations (Figure 6).

**Figure 6.**
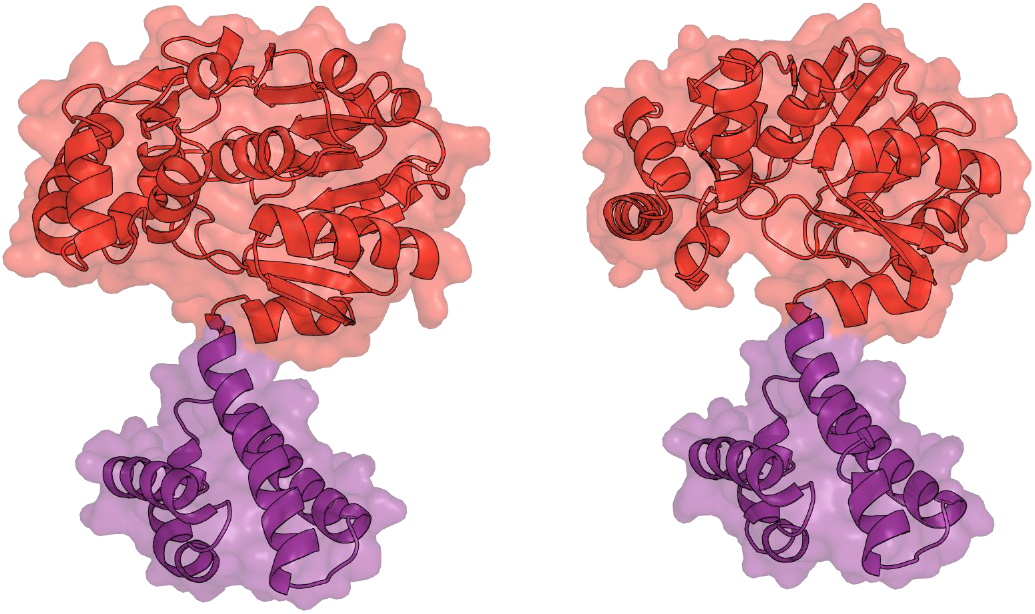
An example of a protein with two open conformations and no closed conformation that we removed from our evaluation set (A6UVT1)

### D.2 Metrics for evaluating switches

We evaluate the designed sequences following a procedure close to that of Abrudan et al. [2026]. Given a designed sequence *s* and the two ground-truth backbones *x*_*c*_ (closed) and *x*_*o*_ (open), we run AlphaFold3 twice: once with *s* and *x*_*c*_ supplied as the template, and once with *s* and *x*_*o*_. Each run samples 15 structures (3 random seeds, 5 samples per seed) as opposed to 5 in the original procedure, from which we extract the predicted backbones 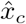 and 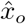 and compare each to its input template.

The comparison is domain-aware: for a template *x* and prediction 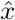 we rigidly align them on each of the two domains in turn and measure the displacement of the *other* domain. RMSD_1_ aligns on the second domain and scores the first; RMSD_2_ aligns on the first domain and scores the second. Their average is a per-state self-consistency score,

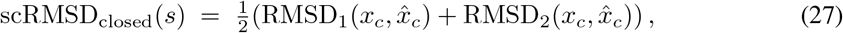

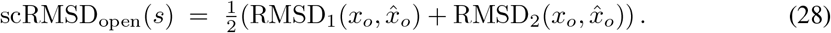

A successful switch must adopt *both x*_*c*_ and *x*_*o*_, so its quality is limited by the worse of the two states:

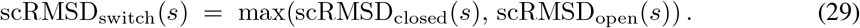

This domain-aligned quantity is the LRMS [Basu and Wallner, 2016], the appropriate measure when the relative arrangement of two parts, here the two domains of a protein, is what matters. In the literature, an LRMS value of 10 Å is treated as the boundary between correct and incorrect predictions [Basu and Wallner, 2016]; we found this too lenient for our setting and use 5 Å throughout as the threshold separating a correct from an incorrect fold.

Each AlphaFold3 run returns 15 predictions, so every sequence has 15 values each of scRMSD_closed_ and scRMSD_open_. Collapsing each set to its single best (minimum) value before combining them discards the spread of the predictions and optimistically overstates a sequence’s switch quality. To retain this distributional information while remaining comparable to Abrudan et al. [2026], we report scRMSD_switch_ as a function of the number of AlphaFold3 predictions considered, *n* (the #AF3 Preds column of Table 1). For a given *n*, we draw *n* of the 15 closed and *n* of the 15 open predictions at random, take the minimum scRMSD within each state, and combine them as above; averaging over 100 independent draws gives the reported value together with an error estimate. Small *n* reflects the switch quality expected from few predictions, whereas *n*=15 uses all predictions and recovers the (deterministic) best-of-all. Unless stated otherwise our figures use *n*=5, and the full dependence on *n* is reported separately.

**Table 1:** Switch self-consistency of designed sequences. scRMSD_switch_ (↓, in Å) and scTM_switch_ (↑), obtained by folding each design with AlphaFold3 against the closed (*x*_*c*_) and open (*x*_*o*_) reference conformations (15 predictions per state) and taking the worse of the two states. Each block reports the metric as the best over a given number of AlphaFold3 predictions per state (#AF3 Preds): for fewer than 15 the predictions are subsampled from the 15 available and the score is averaged over a 100-draw bootstrap, while #AF3 Preds=15 uses all predictions (hence *±* 0). For each protein the single best-scoring design is reported; each entry is the mean over the 20 OC23 switch proteins and *±* is the bootstrap standard deviation. Diff-Switch (multi-state design) is compared against the single-state Baseline for both ProteinMPNN and Caliby inverse folding; the better method is shown in bold.

| Ref. | Metric | #AF3 Preds | ProteinMPNN |  | Caliby |  |
| --- | --- | --- | --- | --- | --- | --- |
|  |  |  | Baseline | Diff-Switch | Baseline | Diff-Switch |
| Closed | $\text{scRMSD}_{\text{switch}} \downarrow$ | 1 | 7.70±0.54 | **5.12**±0.16 | 9.13±0.48 | **6.39**±0.39 |
| | | 5 | 5.86±0.66 | **4.26**±0.09 | 8.00±0.65 | **5.24**±0.63 |
| | | 15 | 4.74±0.00 | **3.88**±0.00 | 6.73±0.00 | **3.91**±0.00 |
| | $\text{scTM}_{\text{switch}} \uparrow$ | 1 | 0.47±0.01 | **0.57**±0.01 | 0.44±0.01 | **0.55**±0.01 |
| | | 5 | 0.55±0.02 | **0.64**±0.01 | 0.48±0.02 | **0.62**±0.01 |
| | | 15 | 0.60±0.00 | **0.67**±0.00 | 0.52±0.00 | **0.67**±0.00 |
| Open | $\text{scRMSD}_{\text{switch}} \downarrow$ | 1 | 7.38±0.41 | **5.63**±0.23 | 8.51±0.21 | **6.41**±0.34 |
| | | 5 | 5.98±0.31 | **4.70**±0.13 | 7.55±0.13 | **4.85**±0.16 |
| | | 15 | 5.07±0.00 | **4.31**±0.00 | 7.11±0.00 | **4.28**±0.00 |
| | $\text{scTM}_{\text{switch}} \uparrow$ | 1 | 0.50±0.02 | **0.59**±0.01 | 0.48±0.01 | **0.55**±0.01 |
| | | 5 | 0.57±0.02 | **0.65**±0.01 | 0.53±0.01 | **0.63**±0.01 |
| | | 15 | 0.62±0.00 | **0.67**±0.00 | 0.57±0.00 | **0.66**±0.00 |

The TM-score analogue is obtained identically, taking the best-of-*n* within each state as the *maximum* (a higher TM-score is better) and the switch value as the worse state, scTM_switch_(*s*) = min(scTM_closed_(*s*), scTM_open_(*s*)); we use 0.5 as its correct/incorrect threshold.

Finally, each protein yields several designed sequences. We report, for each protein, the single design with the best mean scRMSD_switch_ (respectively scTM_switch_) across the bootstrap: a best-average-sequence selection that reflects the design one would actually carry forward. Table 1 summarizes these metrics across the dataset.

### D.3 Inverse Folding Methods

ProteinMPNN [Dauparas et al., 2022] is a message-passing neural network for fixed-backbone inverse folding, widely used for protein sequence design. Given a target backbone, it predicts and samples amino-acid sequences likely to encode that structure. Caliby [Shuai et al., 2025] is an ensemble-conditioned sequence design model that represents sequence compatibility through a neural-network-derived Potts energy function. Unlike single-structure inverse-folding models, Caliby can directly condition on structural ensembles by combining constraints across conformers. We use both ProteinMPNN and Caliby as downstream sequence-design backends to test whether Diff-Switch improves switch design independently of the specific inverse-folding model.

ProteinMPNN designs used temperature 0.1, 0.3 Å backbone noise, and 16 sampled sequences per target condition. Caliby designs used the “caliby” checkpoint with 16 sampled sequences per target condition and ensemble conditioning over the staged conformers. Same noise level (0.3 Å) and temperature (0.1) was also used in Caliby.

### D.4 Further results for 5.1

Table 1 and Figures 7 to 10 are supplementary results for domain-rearrangement evaluation.

**Figure 7.**
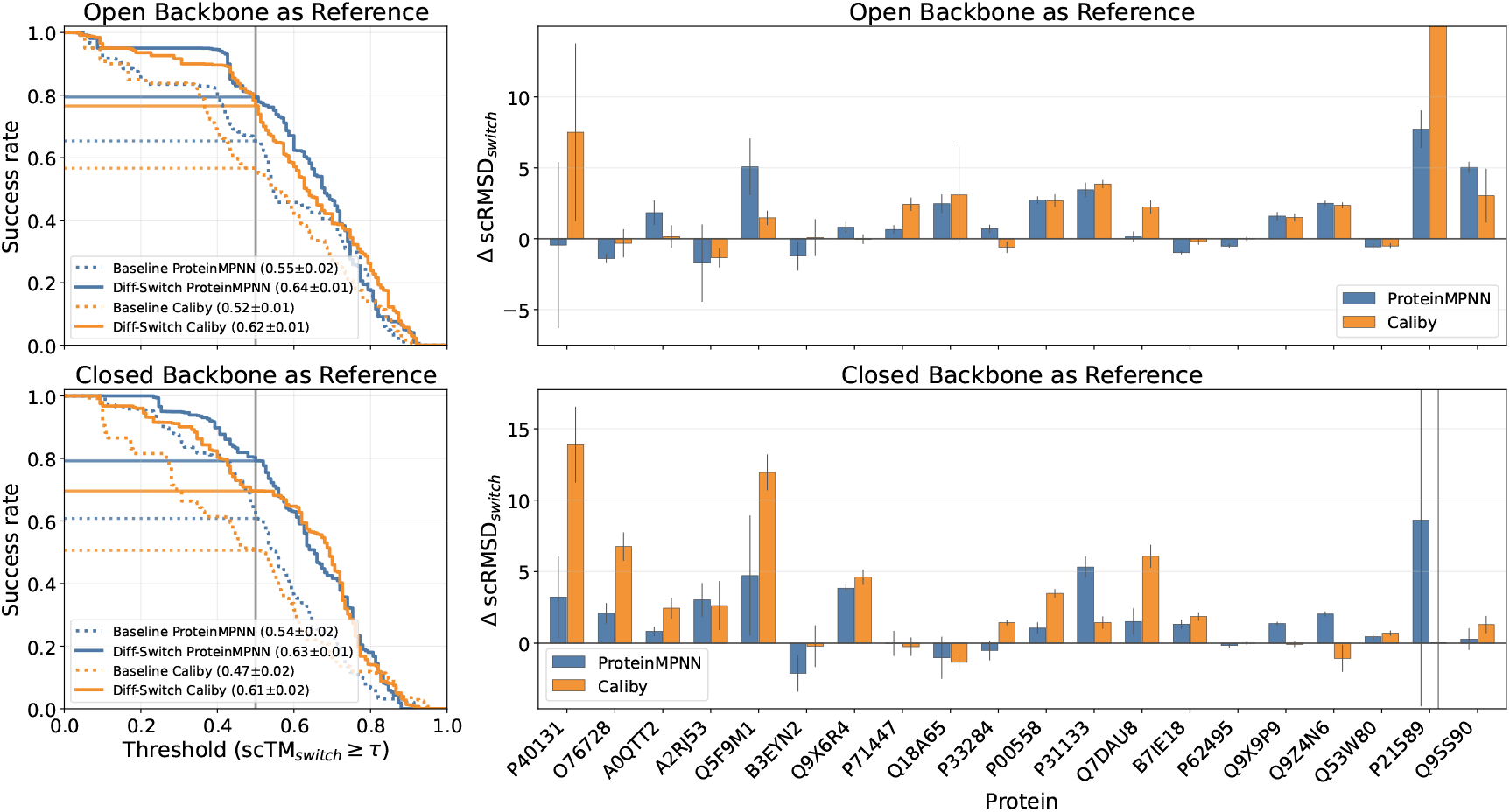
Complementary evaluation of the performance of Diff-Switch on domain-rearrangement protein switches. **Left:** Success rates defined by different scTM_switch_ thresholds *τ* for predicted structures of baseline sequences and those generated for backbones emerging from Diff-Switch. Sequences were generated using either ProteinMPNN or Caliby. The vertical line marks *τ* = 0.5, the value corresponding above which a prediction is considered correct. AUCs are reported in the legend. **Right:** Differences between scRMSD_switch_ scores calculated for structures predicted from sequences that were generated using the Diff-Switch pipeline and the baseline sequences. Positive values indicate better agreement with experimental structures for sequences emerging from the Diff-Switch pipeline. We investigate the noisy case P21589 in Figure 5.

**Figure 8.**
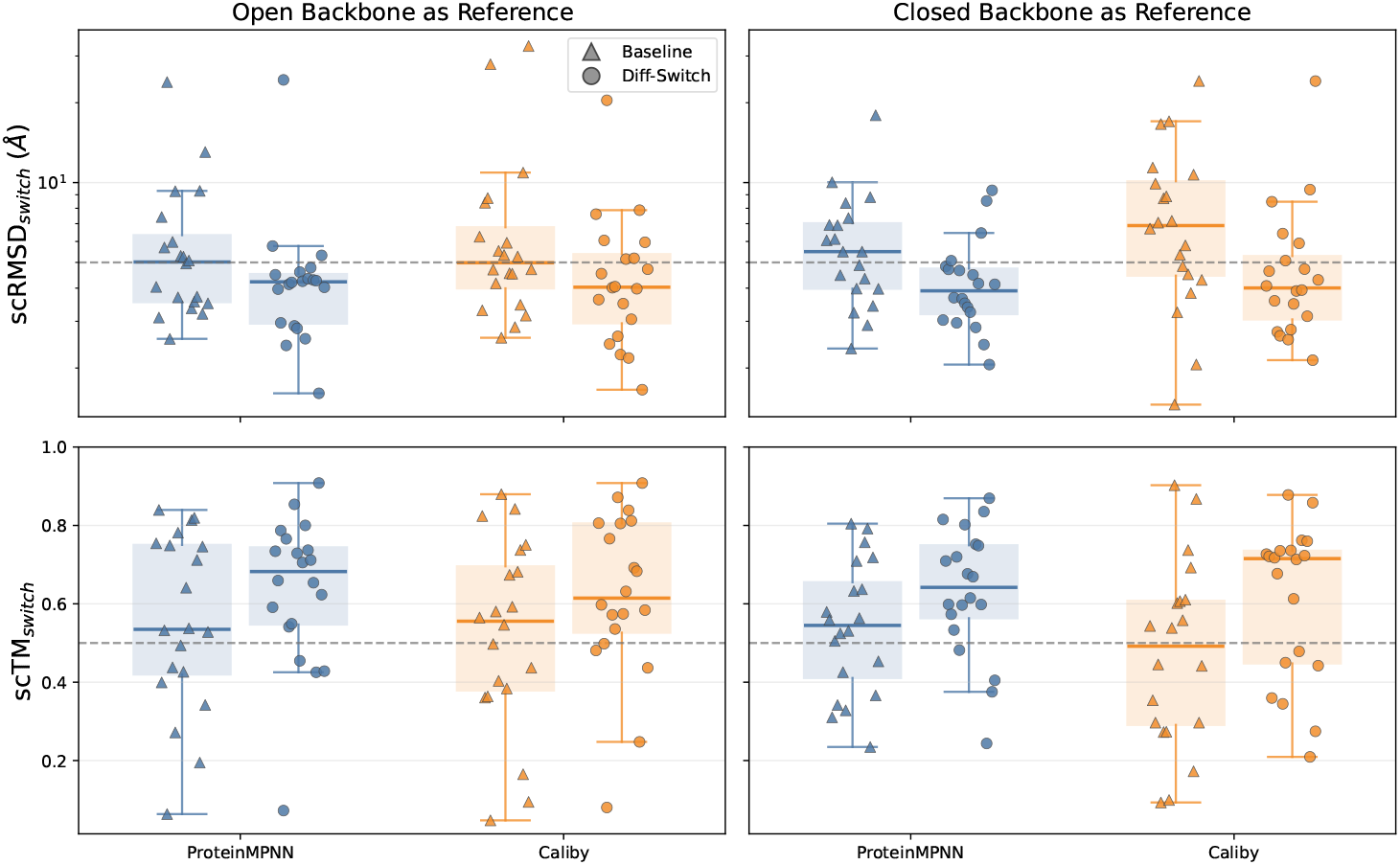
Evaluation of Diff-Switch and baseline designs under domain-aligned switch self-consistency metrics. **Top:** scRMSD_switch_ and **Bottom:** scTM_switch_, with the ground-truth open (left) and closed (right) conformations used as the reference. Within each panel, baseline and Diff-Switch designs are contrasted for ProteinMPNN and Caliby; each point is one protein’s best designed sequence and the boxes summarize the median and inter-quartile range over the switch proteins. The dashed line in the top row marks the 5 Å threshold (see Appendix D).

**Figure 9.**
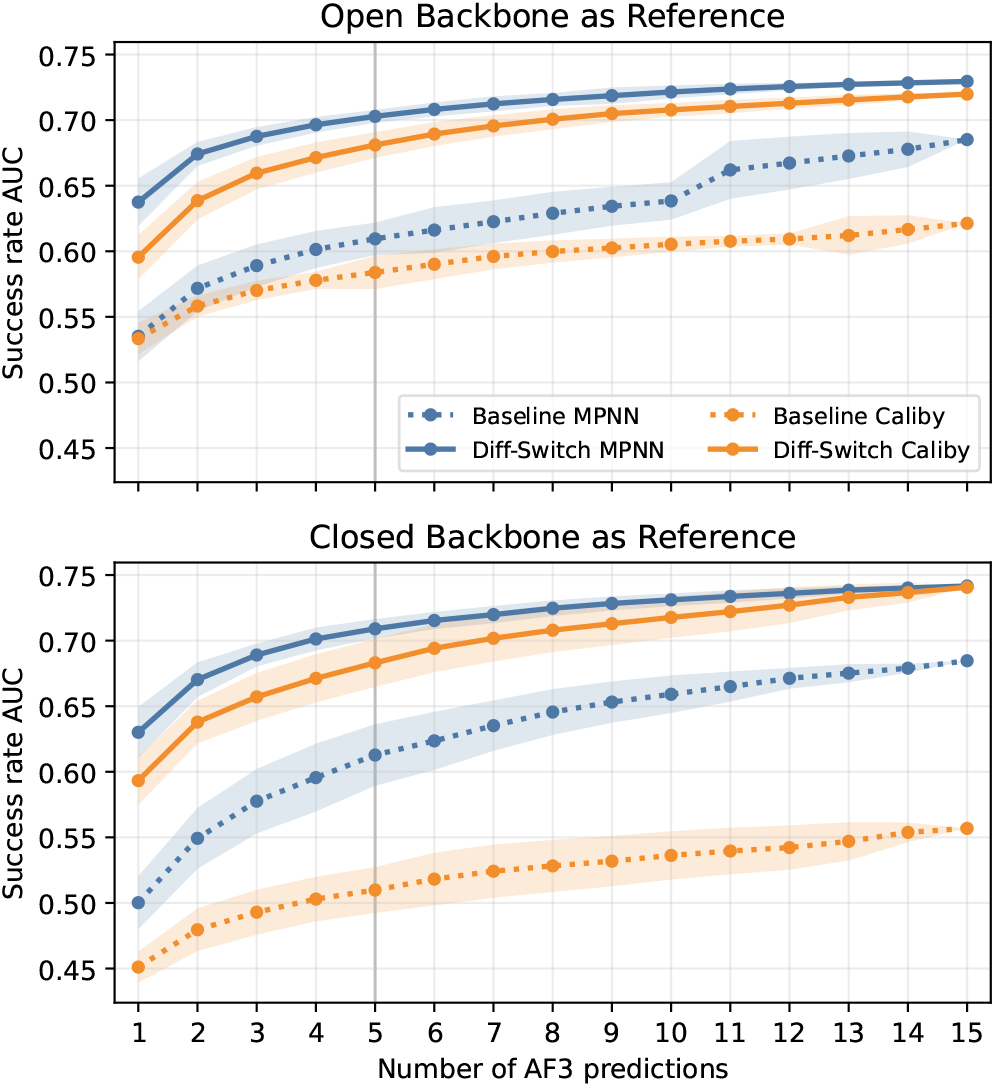
Evaluation of how the Diff-Switch and baseline success rates depend on the AlphaFold3 prediction budget, on domain-rearrangement protein switches. **Top:** closed backbone and **Bottom:** open backbone used as the reference conformation. Each panel plots the area under the scRMSD_switch_ success-rate curve (the fraction of proteins meeting scRMSD_switch_ ≤ *τ*, integrated over thresholds *τ* up to 15 Å) as a function of the number of AlphaFold3 predictions, the number of predictions per state from which the best is retained. Curves are shown for baseline and Diff-Switch sequences generated with either ProteinMPNN or Caliby; the vertical line marks 5 AlphaFold3 predictions and shaded bands denote the bootstrap spread.

**Figure 10.**
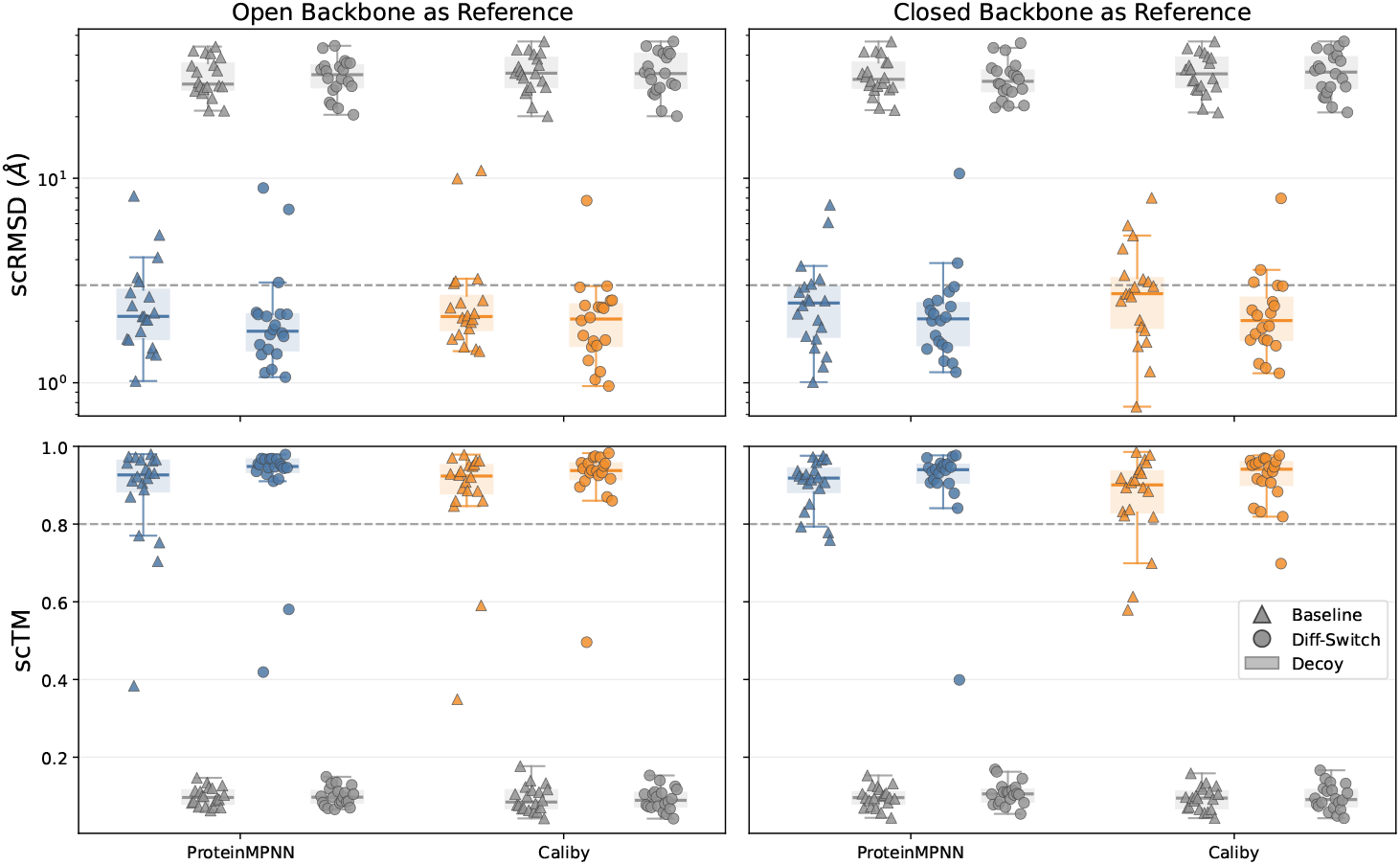
Evaluation of Diff-Switch and baseline designs against a template-swap decoy control, under whole-structure self-consistency metrics, on domain-rearrangement protein switches. **Top:** global scRMSD (all-residue C*α* RMSD) and **Bottom:** global scTM, with the open (left) and closed (right) conformations used as the reference. Within each panel, designs are shown for ProteinMPNN and Caliby, contrasting the Baseline (triangles) and Diff-Switch (circles) pipelines. Each design is paired with its own decoy (grey): a negative control obtained by folding the same designed sequence with AlphaFold3 against a swapped reference template, so that a sequence with no genuine preference for its target backbone would score no better than its decoy. Each point is one protein’s best design, or that design’s decoy, and the boxes summarize the median and inter-quartile range over the switch proteins. Dashed lines mark 3 Å (top) and 0.8 (bottom).

### D.5 Details of phosphorylation-induced protein switch design (section 5.2)

We took a de-novo designed protein with *αβ* structure from PDB (7BPL, Minami et al. [2023]). Residues 1-102 contain four helices and four beta strands. The C-terminus helix (residues 90-102) contains the phosphorylation site RRAS. The rest (residues 1-83) are the scaffold. There is a linker between the scaffold and the helix. The backbone structure of the residues 1-102 acts as the reference backbone *x*_ref_. See Figure 11 for complementary information.

**Figure 11.**
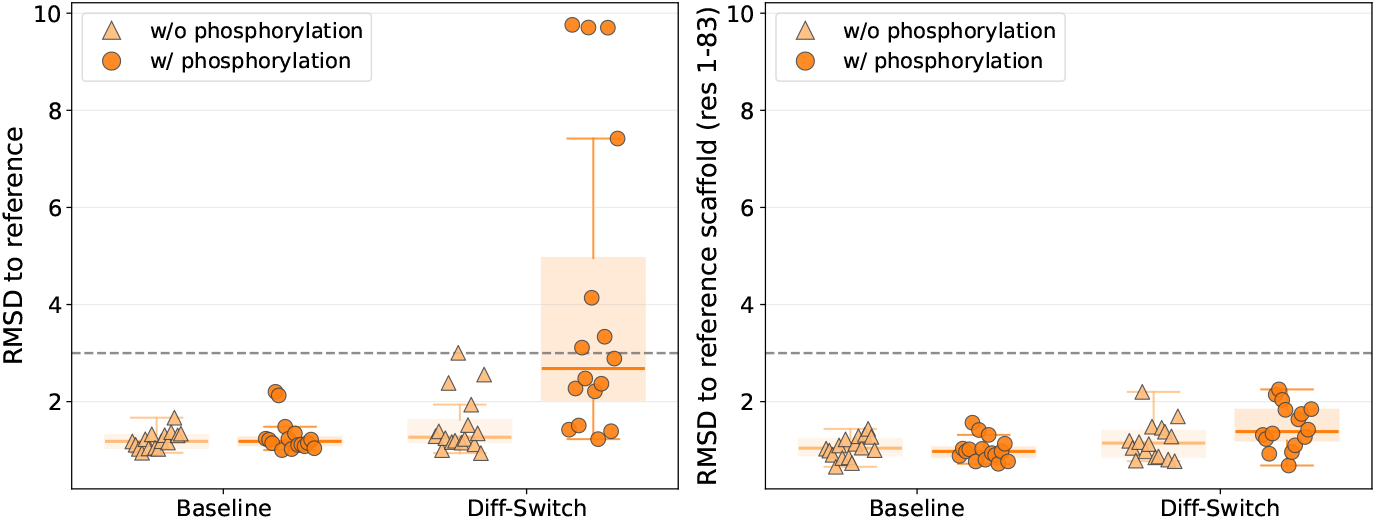
Structural fidelity of Diff-Switch designs to the reference structure in the phosphorylation-induced protein switch experiment. **Left:** Full-structure C*α* RMSD to the reference structure (residues 1-102; global superposition), shown per designed sequence for w/o phosphorylation (unmodified; light orange triangles) and w/ phosphorylation (Residue 95 converted from serine to phosphoserine; dark orange circles). Sequences designed with Diff-Switch are compared with Baseline on the horizontal axis. Diff-Switch sequences are compared with Baseline designs. The dashed line marks a 3 Å cutoff. **Right:** Reference-scaffold C*α* RMSD to the reference scaffold (residues 1-83), shown in the same format.

## E Declaration of Experiments compute resources

The experiments conducted in this paper are not resource-intensive. We used a single 96GB NVIDIA RTX PRO 6000 GPU.

## F Declaration of LLM usage

We used LLMs to assist with coding, manuscript polishing, and visualization of illustrative figures.

## G Licenses for existing assets

Our implementation is based on the following codebases:

- https://github.com/bytedance/PXDesign [Team et al., 2025] (Apache 2.0 License)
- https://github.com/dauparas/ProteinMPNN [Dauparas et al., 2022] (MIT License)
- https://github.com/ProteinDesignLab/caliby [Shuai et al., 2025] (Apache 2.0 License)
- https://github.com/Alex-Abrudan/DynamicMPNN [Abrudan et al., 2026] (MIT License)
- https://github.com/google-deepmind/alphafold3 [Abramson et al., 2024] (CC-BY-NC-SA 4.0 and AlphaFold 3 License)

## H Impact Statement

De novo protein switch design has potential positive applications in synthetic biology, biosensors, and therapeutic engineering, where dynamic responses to molecular stimuli are functionally important and difficult to engineer with conventional approaches. As with any general-purpose protein design method, there is a dual-use concern that improvements in functional design could in principle assist the design of harmful biomolecules. Diff-Switch, however, operates on top of a publicly available pretrained backbone diffusion model and does not unlock new structural capabilities beyond those already accessible through that model; it directs the prior towards switch-compatible ensembles using sampling techniques that have long been standard in the molecular dynamics literature. The marginal biosecurity risk introduced by our method beyond that of the underlying pretrained model is therefore limited, and we do not release any new pretrained model.

## References

J. Abramson, J. Adler, J. Dunger, R. Evans, T. Green, A. Pritzel, O. Ronneberger, L. Willmore, A. J. Ballard, J. Bambrick, S. W. Bodenstein, D. A. Evans, C.-C. Hung, M. O’Neill, D. Reiman, K. Tunya-suvunakool, Z. Wu, A. Žemgulytė, E. Arvaniti, C. Beattie, O. Bertolli, A. Bridgland, A. Cherepanov, M. Congreve, A. I. Cowen-Rivers, A. Cowie, M. Figurnov, F. B. Fuchs, H. Gladman, R. Jain, Y. A. Khan, C. M. R. Low, K. Perlin, A. Potapenko, P. Savy, S. Singh, A. Stecula, A. Thillaisundaram, C. Tong, S. Yakneen, E. D. Zhong, M. Zielinski, A. Žídek, V. Bapst, P. Kohli, M. Jaderberg, D. Hassabis, and J. M. Jumper. Accurate structure prediction of biomolecular interactions with AlphaFold 3. Nature, 630(8016):493–500, June 2024. ISSN 1476-4687. doi: 10.1038/s41586-024-07487-w. URL https://www.nature.com/articles/s41586-024-07487-w.

A. Abrudan, S. P. Ojeda, C. K. Joshi, M. Greenig, F. Engelberger, A. Khmelinskaia, J. Meiler, M. Vendruscolo, and T. P. J. Knowles. MULTI-STATE PROTEIN SEQUENCE DESIGN WITH DYNAMICMPNN. 2026.

R. G. Alberstein, A. B. Guo, and T. Kortemme. Design principles of protein switches. Current Opinion in Structural Biology, 72:71–78, Feb. 2022. ISSN 0959-440X. doi: 10.1016/j.sbi.2021.08.004. URL https://www.sciencedirect.com/science/article/pii/S0959440X21001263.

A. Bansal, H.-M. Chu, A. Schwarzschild, S. Sengupta, M. Goldblum, J. Geiping, and T. Goldstein. Universal guidance for diffusion models. In Proceedings of the IEEE/CVF conference on computer vision and pattern recognition, pages 843–852, 2023.

A. Barducci, G. Bussi, and M. Parrinello. Well-Tempered Metadynamics: A Smoothly Converging and Tunable Free-Energy Method. Physical Review Letters, 100(2):020603, Jan. 2008. ISSN 0031-9007, 1079-7114. doi: 10.1103/PhysRevLett.100.020603. URL https://link.aps.org/doi/10.1103/PhysRevLett.100.020603.

S. Basu and B. Wallner. DockQ: A Quality Measure for Protein-Protein Docking Models. PLOS ONE, 11(8):e0161879. Aug. 2016. ISSN 1932-6203. doi: 10.1371/journal.pone.0161879. URL https://journals.plos.org/plosone/article?id=10.1371/journal.pone.0161879.

N. R. Bennett, B. Coventry, I. Goreshnik, B. Huang, A. Allen, D. Vafeados, Y. P. Peng, J. Dauparas, M. Baek, L. Stewart, F. DiMaio, S. De Munck, S. N. Savvides, and D. Baker. Improving de novo protein binder design with deep learning. Nature Communications, 14(1):2625, May 2023. ISSN 2041-1723. doi: 10.1038/s41467-023-38328-5. URL https://www.nature.com/articles/s41467-023-38328-5.

H. M. Berman, J. Westbrook, Z. Feng, G. Gilliland, T. N. Bhat, H. Weissig, I. N. Shindyalov, and P. E. Bourne. The protein data bank. Nucleic Acids Research, 28(1):235–242, 2000. doi: 10.1093/nar/28.1.235. URL https://doi.org.

A. Bisello, M. Chorev, M. Rosenblatt, L. Monticelli, D. F. Mierke, and S. L. Ferrari. Selective Ligand-induced Stabilization of Active and Desensitized Parathyroid Hormone Type 1 Receptor Conformations. Journal of Biological Chemistry, 277(41):38524–38530, Oct. 2002. ISSN 00219258. doi: 10.1074/jbc.M202544200. URL https://linkinghub.elsevier.com/retrieve/pii/S002192581836335X.

T. J. Brunette, F. Parmeggiani, P.-S. Huang, G. Bhabha, D. C. Ekiert, S. E. Tsutakawa, G. L. Hura, J. A. Tainer, and D. Baker. Exploring the repeat protein universe through computational protein design. Nature, 528(7583):580–584, Dec. 2015. ISSN 1476-4687. doi: 10.1038/nature16162. URL https://www.nature.com/articles/nature16162.

S. Buckley, Y. Miao, M. Idris, P.-W. Lee, L. Scheller, R. Riek, S. J. Maerkl, L. A. Abriata, and B. E. Correia. De novo design of phosphorylation-induced protein switches for synthetic signaling in cells, Sept. 2025. URL https://www.biorxiv.org/content/10.1101/2025.09.10.675034v1. ISSN: 2692-8205 Pages: 2025.09.10.675034 Section: New Results.

H. Chung, J. Kim, M. T. Mccann, M. L. Klasky, and J. C. Ye. Diffusion posterior sampling for general noisy inverse problems. arXiv preprint arXiv:2209.14687, 2022.

J. Dauparas, I. Anishchenko, N. Bennett, H. Bai, R. J. Ragotte, L. F. Milles, B. I. M. Wicky Courbet, R. J. de Haas, N. Bethel, P. J. Y. Leung, T. F. Huddy, S. Pellock, D. Tischer, F. Chan Koepnick, H. Nguyen, A. Kang, B. Sankaran, A. K. Bera, N. P. King, and D. Baker. Robust deep learning–based protein sequence design using ProteinMPNN. Science, 378(6615):49–56, Oct. 2022. doi: 10.1126/science.add2187. URL https://www.science.org/doi/10.1126/science.add2187.

P. Del Moral, A. Doucet, and A. Jasra. Sequential Monte Carlo samplers. Journal of the Royal Statistical Society: Series B (Statistical Methodology), 68(3):411–436, 2006. doi: 10.1111/j.1467-9868.2006.00553.x.

P. Dhariwal and A. Nichol. Diffusion models beat GANs on image synthesis. In Advances in Neural Information Processing Systems, volume 34, pages 8780–8794, 2021.

C. Discovery, J. Boitreaud, J. Dent, M. McPartlon, J. Meier, V. Reis, A. Rogozhnikov, and K. Wu. Chai-1: Decoding the molecular interactions of life, Oct. 2024. URL https://www.biorxiv.org/content/10.1101/2024.10.10.615955v2. Pages: 2024.10.10.615955 Section: New Results.

T. Geffner, K. Didi, Z. Zhang, D. Reidenbach, Z. Cao, J. Yim, M. Geiger, C. Dallago, E. Kucukbenli, A. Vahdat, and K. Kreis. Proteina: Scaling flow-based protein structure generative models. In International Conference on Learning Representations, 2025.

J. He, J. M. Hernández-Lobato, Y. Du, and F. Vargas. Rne: plug-and-play diffusion inference-time control and energy-based training. arXiv preprint arXiv:2506.05668, 2025.

J. Ho, A. Jain, and P. Abbeel. Denoising diffusion probabilistic models. In Advances in Neural Information Processing Systems, volume 33, pages 6840–6851, 2020.

C. Hsu, R. Verkuil, J. Liu, Z. Lin, B. Hie, T. Sercu, A. Lerer, and A. Rives. Learning inverse folding from millions of predicted structures. In Proceedings of the 39th International Conference on Machine Learning, pages 8946–8970. PMLR, June 2022. URL https://proceedings.mlr.press/v162/hsu22a.html.

J. Hénin, T. Lelièvre, M. R. Shirts, O. Valsson, and L. Delemotte. Enhanced Sampling Methods for Molecular Dynamics Simulations [Article v1.0]. Living Journal of Computational Molecular Science, 4(1):1583–1583, Dec. 2022. ISSN 2575-6524. doi: 10.33011/livecoms.4.1.1583. URL https://livecomsjournal.org/index.php/livecoms/article/view/v4i1e1583.

J. B. Ingraham, M. Baranov, Z. Costello, K. W. Barber, W. Wang, A. Ismail, V. Frappier, D. M. Lord, C. Ng-Thow-Hing, E. R. Van Vlack, S. Tie, V. Xue, S. C. Cowles, A. Leung, J. V. Rodrigues, C. L. Morales-Perez, A. M. Ayoub, R. Green, K. Puentes, F. Oplinger, N. V. Panwar, F. Obermeyer, A. R. Root, A. L. Beam, F. J. Poelwijk, and G. Grigoryan. Illuminating protein space with a programmable generative model. Nature, 623(7989):1070–1078, Nov. 2023. ISSN 1476-4687. doi: 10.1038/s41586-023-06728-8. URL https://www.nature.com/articles/s41586-023-06728-8.

M. Invernizzi and M. Parrinello. Rethinking Metadynamics: From Bias Potentials to Probability Distributions. The Journal of Physical Chemistry Letters, 11(7):2731–2736, Apr. 2020. doi: 10.1021/acs.jpclett.0c00497. URL https://doi.org/10.1021/acs.jpclett.0c00497.

J. Jumper, R. Evans, A. Pritzel, T. Green, M. Figurnov, O. Ronneberger, K. Tunyasuvunakool, R. Bates, A. Žídek, A. Potapenko, A. Bridgland, C. Meyer, S. A. A. Kohl, A. J. Ballard, A. Cowie, B. Romera-Paredes, S. Nikolov, R. Jain, J. Adler, T. Back, S. Petersen, D. Reiman, E. Clancy, M. Zielinski, M. Steinegger, M. Pacholska, T. Berghammer, S. Bodenstein, D. Silver, O. Vinyals, A. W. Senior, K. Kavukcuoglu, P. Kohli, and D. Hassabis. Highly accurate protein structure prediction with AlphaFold. Nature, 596(7873):583–589, Aug. 2021. ISSN 1476-4687. doi: 10.1038/s41586-021-03819-2. URL https://www.nature.com/articles/s41586-021-03819-2.

Y. Kalakoti and B. Wallner. AFsample2 predicts multiple conformations and ensembles with AlphaFold2. Communications Biology, 8(1):373, Mar. 2025. ISSN 2399-3642. doi: 10.1038/s42003-025-07791-9. URL https://www.nature.com/articles/s42003-025-07791-9.

T. Karras, M. Aittala, T. Aila, and S. Laine. Elucidating the design space of diffusion-based generative models. In Advances in Neural Information Processing Systems, volume 35, 2022.

A. Laio and M. Parrinello. Escaping free-energy minima. Proceedings of the National Academy of Sciences of the United States of America, 99(20):12562–12566, Oct. 2002. ISSN 0027-8424. doi: 10.1073/pnas.202427399. URL https://pmc.ncbi.nlm.nih.gov/articles/PMC130499/.

H. Y. I. Lam, S. P. Ojeda, M. Brezinova, J. Hanke, X. E. Ong, Y. Mu, and M. Vendruscolo. Metadiffusion: inference-time meta-energy biasing of biomolecular diffusion models, Feb. 2026. URL https://www.biorxiv.org/content/10.64898/2026.02.10.704873v1. ISSN: 2692-8205 Pages: 2026.02.10.704873 Section: New Results.

R. A. Langan, S. E. Boyken, A. H. Ng, J. A. Samson, G. Dods, A. M. Westbrook, T. H. Nguyen, M. J. Lajoie, Z. Chen, S. Berger, V. K. Mulligan, J. E. Dueber, W. R. P. Novak, H. El-Samad, and D. Baker. De novo design of bioactive protein switches. Nature, 572(7768):205–210, Aug. 2019. ISSN 1476-4687. doi: 10.1038/s41586-019-1432-8. URL https://www.nature.com/articles/s41586-019-1432-8.

A. Leaver-Fay, M. Tyka, S. M. Lewis, O. F. Lange, J. Thompson, R. Jacak, K. Kaufman, P. D. Renfrew, C. A. Smith, W. Sheffler, I. W. Davis, S. Cooper, A. Treuille, D. J. Mandell, F. Richter, Y.-E. A. Ban, S. J. Fleishman, J. E. Corn, D. E. Kim, S. Lyskov, M. Berrondo, S. Mentzer, Z. Popović, J. J. Havranek, J. Karanicolas, R. Das, J. Meiler, T. Kortemme, J. J. Gray, B. Kuhlman, D. Baker, and P. Bradley. ROSETTA3: an object-oriented software suite for the simulation and design of macromolecules. Methods in Enzymology, 487:545–574, 2011. ISSN 1557-7988. doi: 10.1016/B978-0-12-381270-4.00019-6.

Y. Lin, M. Lee, Z. Zhang, and M. AlQuraishi. Out of many, one: Designing and scaffolding proteins at the scale of the structural universe with Genie 2. arXiv preprint arXiv:2405.15489, 2024.

S. Minami, N. Kobayashi, T. Sugiki, T. Nagashima, T. Fujiwara, R. Tatsumi-Koga, G. Chikenji, and N. Koga. Exploration of novel αβ-protein folds through de novo design. Nature Structural & Molecular Biology, 30(8):1132–1140, Aug. 2023. ISSN 1545-9985. doi: 10.1038/s41594-023-01029-0. URL https://www.nature.com/articles/s41594-023-01029-0.

J. Nam, B. Máté, A. P. Toshev, M. Kaniselvan, R. Gómez-Bombarelli, R. T. Chen, B. Wood, G.-H. Liu, and B. K. Miller. Enhancing diffusion-based sampling with molecular collective variables. arXiv preprint arXiv:2510.11923, 2025.

J. Ohnuki and K.-i. Okazaki. Enhanced sampling of protein conformations in AlphaFold3 with repulsive bias in the diffusion generative model, Dec. 2025. URL https://www.biorxiv.org/content/10.64898/2025.12.17.693105v1. ISSN: 2692-8205 Pages: 2025.12.17.693105 Section: New Results.

S. Passaro, G. Corso, J. Wohlwend, M. Reveiz, S. Thaler, V. R. Somnath, N. Getz, T. Portnoi, J. Roy, H. Stark, D. Kwabi-Addo, D. Beaini, T. Jaakkola, and R. Barzilay. Boltz-2: Towards Accurate and Efficient Binding Affinity Prediction, June 2025. URL https://www.biorxiv.org/content/10.1101/2025.06.14.659707v1. Pages: 2025.06.14.659707 Section: New Results.

F. Praetorius, P. J. Y. Leung, M. H. Tessmer, A. Broerman, C. Demakis, A. F. Dishman, A. Pillai, A. Idris, D. Juergens, J. Dauparas, X. Li, P. M. Levine, M. Lamb, R. K. Ballard, S. R. Gerben, H. Nguyen, A. Kang, B. Sankaran, A. K. Bera, B. F. Volkman, J. Nivala, S. Stoll, and D. Baker. Design of stimulus-responsive two-state hinge proteins. Science, 381(6659):754–760, Aug. 2023. doi: 10.1126/science.adg7731. URL https://www.science.org/doi/10.1126/science.adg7731.

D. D. Richman, J. Karaguesian, C.-M. Suomivuori, and R. O. Dror. Unlocking hidden biomolecular conformational landscapes in diffusion models at inference time, Feb. 2026. URL http://arxiv.org/abs/2512.03312. arXiv:2512.03312 [q-bio].

R. W. Shuai, T. Lu, S. Bhatti, P. Kouba, and P.-S. Huang. Ensemble-conditioned protein sequence design with Caliby, Oct. 2025. URL https://www.biorxiv.org/content/10.1101/2025.09.30.679633v4. ISSN: 2692-8205 Pages: 2025.09.30.679633 Section: New Results.

R. Singhal, Z. Horvitz, R. Teehan, M. Ren, Z. Yu, K. McKeown, and R. Ranganath. A general framework for inference-time scaling and steering of diffusion models. arXiv preprint arXiv:2501.06848, 2025.

M. Skreta, T. Akhound-Sadegh, V. Ohanesian, R. Bondesan, A. Aspuru-Guzik, A. Doucet, R. Brekelmans, A. Tong, and K. Neklyudov. Feynman-kac correctors in diffusion: Annealing, guidance, and product of experts. arXiv preprint arXiv:2503.02819, 2025.

Y. Song, J. Sohl-Dickstein, D. P. Kingma, A. Kumar, S. Ermon, and B. Poole. Score-based generative modeling through stochastic differential equations. In International Conference on Learning Representations, 2021.

H. Stark, F. Faltings, M. Choi, Y. Xie, E. Hur, T. O’Donnell, A. Bushuiev, T. Uçar, S. Passaro, W. Mao, M. Reveiz, R. Bushuiev, T. Pluskal, J. Sivic, K. Kreis, A. Vahdat, S. Ray, J. T. Goldstein, A. Savinov, J. A. Hambalek, A. Gupta, D. A. Taquiri-Diaz, Y. Zhang, A. K. Hatstat, A. Arada, N. H. Kim, E. Tackie-Yarboi, D. Boselli, L. Schnaider, C. C. Liu, G.-W. Li, D. Hnisz, D. M. Sabatini, W. F. DeGrado, J. Wohlwend, G. Corso, R. Barzilay, and T. Jaakkola. BoltzGen: Toward Universal Binder Design, Nov. 2025. URL https://www.biorxiv.org/content/10.1101/2025.11.20.689494v1. ISSN: 2692-8205 Pages: 2025.11.20.689494 Section: New Results.

J. B. Stiller, R. Otten, D. Häussinger, P. S. Rieder, D. L. Theobald, and D. Kern. Structure determination of high-energy states in a dynamic protein ensemble. Nature, 603(7901): 528–535, Mar. 2022. ISSN 0028-0836. doi: 10.1038/s41586-022-04468-9. URL https://pmc.ncbi.nlm.nih.gov/articles/PMC9126080/.

P. Team, M. Ren, J. Sun, J. Guan, C. Liu, C. Gong, Y. Wang, L. Wang, Q. Cai, W. Ma, Y. Zhang, Z. Liu, H. Zhang, X. Chen, and W. Xiao. PXDesign: Fast, Modular, and Accurate De Novo Design of Protein Binders, Dec. 2025. URL https://www.biorxiv.org/content/10.1101/2025.08.15.670450v3. ISSN: 2692-8205 Pages: 2025.08.15.670450 Section: New Results.

P. Team, Y. Zhang, C. Gong, H. Zhang, W. Ma, Z. Liu, X. Chen, J. Guan, L. Wang, Y. Yang, Y. Xia, and W. Xiao. Protenix-v1: Toward High-Accuracy Open-Source Biomolecular Structure Prediction, Feb. 2026. URL https://www.biorxiv.org/content/10.64898/2026.02.05.703733v3. ISSN: 2692-8205 Pages: 2026.02.05.703733 Section: New Results.

J. L. Watson, D. Juergens, N. R. Bennett, B. L. Trippe, J. Yim, H. E. Eisenach, W. Ahern, A. J. Borst, R. J. Ragotte, L. F. Milles, B. I. M. Wicky, N. Hanikel, S. J. Pellock, A. Courbet, W. Sheffler, J. Wang, P. Venkatesh, I. Sappington, S. V. Torres, A. Lauko, V. De Bortoli, E. Mathieu, S. Ovchinnikov, R. Barzilay, T. S. Jaakkola, F. DiMaio, M. Baek, and D. Baker. De novo design of protein structure and function with RFdiffusion. Nature, 620(7976):1089–1100, Aug. 2023. ISSN 1476-4687. doi: 10.1038/s41586-023-06415-8. URL https://www.nature.com/articles/s41586-023-06415-8.

B. I. M. Wicky, L. F. Milles, A. Courbet, R. J. Ragotte, J. Dauparas, E. Kinfu, S. Tipps, R. D. Kibler, M. Baek, F. DiMaio, X. Li, L. Carter, A. Kang, H. Nguyen, A. K. Bera, and D. Baker. Hallucinating symmetric protein assemblies. Science, 378(6615):56–61, Oct. 2022. doi: 10.1126/science.add1964. URL https://www.science.org/doi/full/10.1126/science.add1964.

K. A. Woll, O. Haji-Ghassemi, and F. Van Petegem. Pathological conformations of disease mutant Ryanodine Receptors revealed by cryo-EM. Nature Communications, 12(1):807, Feb. 2021. ISSN 2041-1723. doi: 10.1038/s41467-021-21141-3. URL https://www.nature.com/articles/s41467-021-21141-3.

L. Wu, B. Trippe, C. Naesseth, D. Blei, and J. P. Cunningham. Practical and asymptotically exact conditional sampling in diffusion models. Advances in Neural Information Processing Systems, 36:31372–31403, 2023.

Y. Xie, L. Winkler, L. Sun, S. Lewis, A. E. Foster, J. J. Luna, T. Hempel, M. Gastegger, Y. Chen, I. Zaporozhets, C. Clementi, C. M. Bishop, and F. Noé. Enhanced Diffusion Sampling: Efficient Rare Event Sampling and Free Energy Calculation with Diffusion Models, Feb. 2026. URL http://arxiv.org/abs/2602.16634. arXiv:2602.16634 [stat] version: 1.

W. Yang, S. Wang, G. R. Lee, J. Z. Zhang, A. Courbet, D. Juergens, X. Wang, T. Schlichthaerle, M. Abedi, R. Ragotte, L. An, I. Kalvet, S. Pellock, L. Mihaljevic, C. Glasscock, A. Pillai, A. Broerman, N. Ennist, E. Haefner, N. McNamara-Bordewick, I. Haydon, L. Stewart, G. Bhardwaj, and D. Baker. The past, present and future of de novo protein design. Nature, 652(8112): 1139–1152, Apr. 2026. ISSN 1476-4687. doi: 10.1038/s41586-026-10328-7. URL https://www.nature.com/articles/s41586-026-10328-7.

J. Yim, B. L. Trippe, V. De Bortoli, E. Mathieu, A. Doucet, R. Barzilay, and T. Jaakkola. SE(3) diffusion model with application to protein backbone generation. In Proceedings of the 40th International Conference on Machine Learning, pages 40001–40039, 2023.

V. Zambaldi, D. La, A. E. Chu, H. Patani, A. E. Danson, T. O. C. Kwan, T. Frerix, R. G. Schneider, D. Saxton, A. Thillaisundaram, Z. Wu, I. Moraes, O. Lange, E. Papa, G. Stanton, V. Martin, S. Singh, L. H. Wong, R. Bates, S. A. Kohl, J. Abramson, A. W. Senior, Y. Alguel, M. Y. Wu, I. M. Aspalter, K. Bentley, D. L. V. Bauer, P. Cherepanov, D. Hassabis, P. Kohli, R. Fergus, and J. Wang. De novo design of high-affinity protein binders with AlphaProteo, Sept. 2024. URL http://arxiv.org/abs/2409.08022. arXiv:2409.08022 [q-bio].

